# Head-on and co-directional RNA polymerase collisions orchestrate bidirectional transcription termination

**DOI:** 10.1101/2022.10.23.513370

**Authors:** Ling Wang, John W. Watters, Xiangwu Ju, Shixin Liu

## Abstract

Genomic DNA is a crowded track where translocating motor proteins frequently collide. It remains unclear whether these collisions, generally thought to occur inadvertently, carry any physiological function. In this work, we developed a single-molecule assay to directly visualize the trafficking of individual *E. coli* RNA polymerases (RNAPs) on DNA. This assay enabled us to test the hypothesis that RNAP collisions drive bidirectional transcription termination of convergent gene pairs. We showed that the head-on collision between two converging RNAPs is necessary to prevent transcriptional readthrough, but insufficient to release the collided RNAPs from the DNA. Remarkably, co-directional collision from a trailing RNAP into the head-on collided complex dramatically increases the termination efficiency. Furthermore, a stem-loop structure formed in the nascent RNA is required for collisions to occur at well-defined positions between gene boundaries. These findings, corroborated by transcriptomic data, establish programmed RNAP collisions as an effective strategy to achieve precise gene expression and imply a broader role of genomic conflicts in cell physiology.

## INTRODUCTION

Efficient and precise termination of transcription ensures the generation of accurate gene products (Porrua et al., 2016; Santangelo and Artsimovitch, 2011). By restricting the movement of RNA polymerases (RNAPs) within specific genomic regions, transcription termination also minimizes the conflicts between RNAPs and other molecular machines traveling on DNA, such as the replication machinery (Garcia-Muse and Aguilera, 2016). In bacteria, two well-known mechanisms have been documented to release the RNAP and nascent RNA at the 3’ end of transcription units. The first mechanism is mediated by an RNA stem-loop structure formed in the nascent transcript (intrinsic termination); the second mechanism relies on an ATP-dependent RNA translocase Rho (Rho-dependent termination) [reviewed in ((Porrua *et al.*, 2016; Ray-Soni et al., 2016; Roberts, 2019)].

Recently, using a transcriptomic method termed SEnd-seq to map full-length bacterial RNAs, we identified a prevalent class of bidirectional transcription terminators located between convergent gene pairs in the *Escherichia coli* genome (Ju et al., 2019). These bidirectional terminators—controlling hundreds of *E. coli* genes—do not resemble intrinsic terminators in the RNA sequence that they encode, nor do they depend on the activity of Rho. Thus, we propose a new model for bacterial transcription termination, in which the physical collisions between convergent transcription elongation complexes (ECs) drive their own dissociation. The model, if proven, could represent a conserved mechanism to terminate gene transcription, given the convergent gene arrangement universally found in nearly all organisms (Callen et al., 2004; Georg and Hess, 2018; Gullerova and Proudfoot, 2012; Hobson et al., 2012; Prescott and Proudfoot, 2002; Shearwin et al., 2005; Yelin et al., 2003). However, direct evidence for such collision-driven termination mechanism is still lacking.

Single-molecule techniques are powerful tools to study dynamic biomolecular processes and have greatly aided our understanding of the mechanism of transcription and its regulation [reviewed in (Bai et al., 2006; Dangkulwanich et al., 2014; Larson et al., 2011; Lee and Myong, 2021; Wang and Greene, 2011)]. The two major classes of single-molecule approaches provide complementary information: fluorescence-based detection reveals the composition of transcription complexes [e.g. (Friedman and Gelles, 2012)], while force-based manipulation reports the position of RNAP on a DNA template [e.g. (Abbondanzieri et al., 2005)]. In this work, we sought to combine these two approaches to visualize the dynamics and outcome of RNAP trafficking and collisions, delineating the molecular events occurring at a bidirectional terminator during convergent transcription.

## RESULTS

### A single-molecule platform for studying RNAP trafficking

We first constructed a DNA template (*Template1*) composed of a transcription unit flanked by biotinylated DNA handles for bead conjugation (**Figure 1A**). The transcription unit contains— from upstream to downstream—a σ^70^-dependent T7A1 promoter, a C-less segment, a cassette harboring seven tandem repeats, a 1,831-basepair (bp) transcribed region, and a λ tR’ intrinsic terminator (**Figure 1A** and **Table S1**). Transcription was initiated and stalled in a test tube by mixing Cy3-labeled *E. coli* RNAP/σ^70^ holoenzyme, *Template1*, and ribonucleotides except CTP. Stalled ECs were injected into the microfluidic flow cell of a dual optical trap instrument combined with multicolor confocal fluorescence microscopy (Candelli et al., 2011; Wasserman et al., 2019). A single DNA template containing the stalled EC was tethered between a pair of streptavidin-coated beads held by optical traps and verified by 2D scanning of the Cy3 fluorescence (**Figure 1B**). The tethered complex was then moved to a separate channel containing the full set of NTPs where unidirectional movement of Cy3-RNAP along the DNA template towards the terminator was observed, indicating successful restart of the stalled EC (**Figure 1C**). Time-resolved positions of the Cy3-RNAP on DNA were converted to distance traveled in base pairs (bp) (**Figure 1D**). The average elongation speed of RNAP derived from these trajectories is 16 ± 6 bp/s (**Figure 1E**, *Inset*), consistent with previous single-molecule measurements under similar conditions (Schafer et al., 1991; Wang et al., 1998).

**Figure 1.**
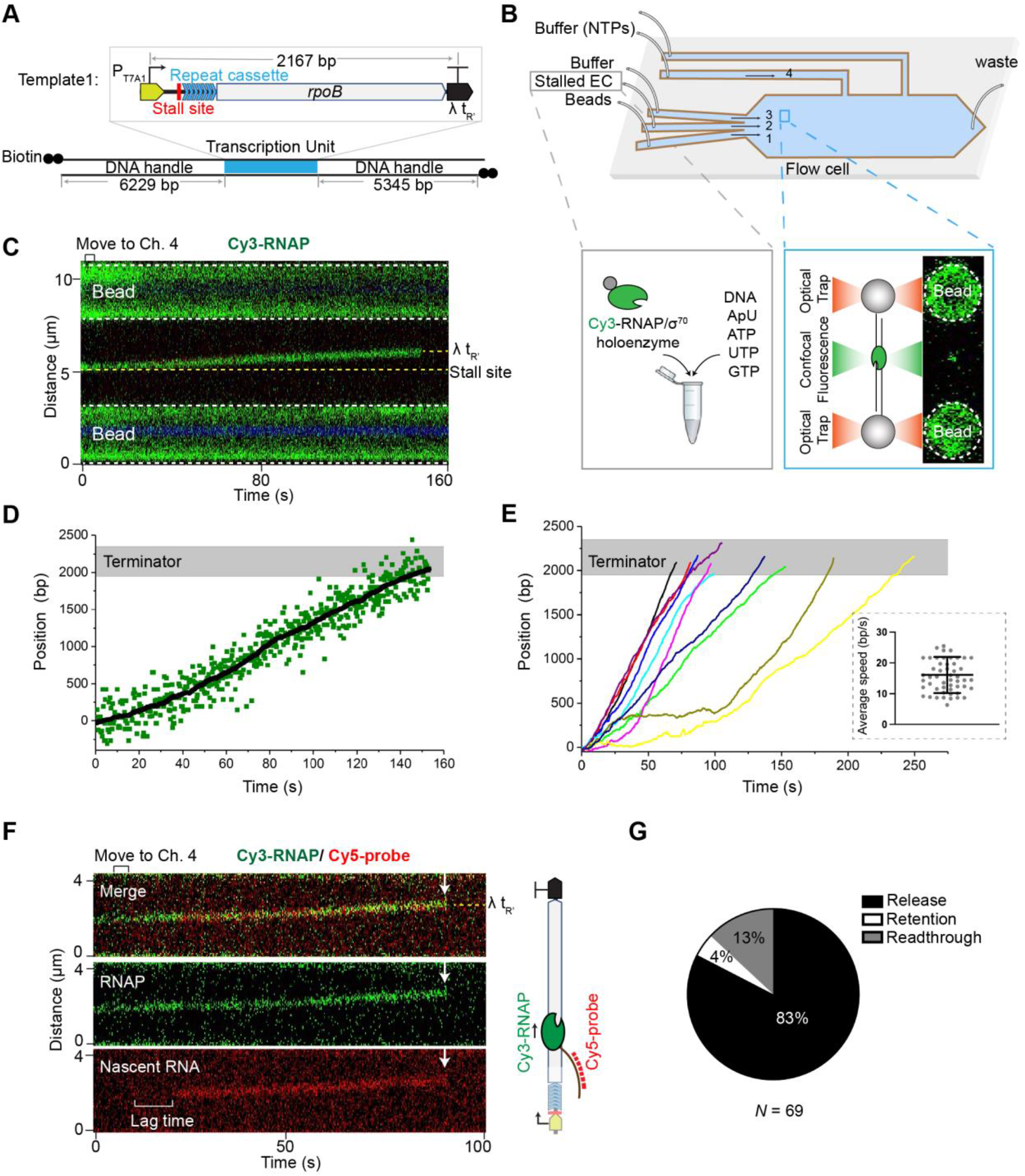
Single-molecule platform for visualizing transcription elongation and termination. (**A**) Schematic of *Template1* that contains a λ tR’ intrinsic terminator. See main text for a detailed description of the transcription unit. (**B**) Single-molecule experimental setup. Streptavidin-coated beads, stalled elongation complexes (ECs) and imaging buffer were flown into channels 1—3, respectively. A single DNA tether loaded with a stalled EC (visualized by Cy3-RNAP fluorescence) was moved to channel 4 to resume transcription. (**C**) A representative kymograph showing unidirectional translocation of a restarted EC along *Template1*. (**D**) Position of the Cy3-RNAP on DNA as a function of time extracted from the kymograph in (C). Raw data and smoothed trajectory ((± 10-s moving average of tracked points) are shown in green dots and black lines, respectively. Gray region indicates the intrinsic terminator position (2150 ± 200 bp). (**E**) Multiple overlaid individual RNAP trajectories on *Template1* show RNAP release within the termination zone. (*Inset*) Distribution of the average elongation speed for 57 individual RNAPs. Error bars represent SD. (**F**) A representative two-color kymograph showing the concomitant translocation and release (white arrows) of both RNAP (green) and nascent RNA (red). (**G**) Pie chart showing the fraction of EC release, retention and readthrough events observed on *Template1*. *N* denotes the total number of ECs. See also Figures S1 and S2.

We found that the trajectories often lost the Cy3 fluorescence signal within the termination zone (**Figure 1E**), with its center corresponding to the terminator position on *Template1* and its width corresponding to the spatial resolution of the instrument. The signal loss was unlikely due to dye photobleaching because the fluorescence lifetime of Cy3-RNAP was much longer under our imaging conditions (>400 s). Thus, the disappearance of Cy3 signal most likely represents release of the RNAP at the intrinsic terminator. To confirm that these translocating ECs were active in synthesizing nascent RNA, we included a Cy5-labeled DNA probe complementary to the RNA sequence transcribed from the repeat cassette (**Figure 1A**). Indeed, we observed the onset of Cy5 signal shortly after the stalled EC resumed translocation (**Figure 1F**). The lag period (22 ± 9 s) accounts for the time needed for the RNAP to transcribe from the stall site to the repeat cassette. The Cy3 and Cy5 signals co-migrated and concomitantly disappeared at the terminator (**Figure 1F** and **Figure S1A**), indicating bona fide transcription termination events at which both RNAP and nascent RNA dissociated from the DNA template. Immediate release of RNAP and RNA was observed in the vast majority (83%) of ECs that reached the terminator site (**Figure 1G**), consistent with the high termination efficiency previously reported for the λ tR’ intrinsic terminator (Rees et al., 1997). We also observed minor subpopulations of ECs that remained associated at the terminator position, as well as those that translocated across the terminator (“*Retention*” and “*Readthrough*” in **Figure 1G** and **Figure S1B-S1C**). These observations highlight the stochastic nature of the termination process and showcase the power of single-molecule analysis in revealing heterogeneous molecular behaviors within a single population. We note that there also existed a fraction of stalled ECs that failed to restart, or restarted but never reached the terminator site (**Figure S2**). In the following we focused on the processive ECs that successfully arrived at the terminator.

### Visualizing collisions between convergent RNAPs

Having demonstrated the utility of our assay in visualizing transcription elongation and termination, we sought to test our hypothesis that collisions between a pair of convergent ECs can drive bidirectional transcription termination. To this end, we constructed a DNA template that supports convergent transcription (*Template2*) (**Figure 2A** and **Table S1**). This template contains two opposite T7A1 promoters, each followed by a transcribed region with a stall site and a distinct repeat cassette, and a bidirectional terminator from the *E. coli* genome located between the *ynaJ-uspE* gene pair (**Figure S3**). We also constructed two unidirectional transcription templates (*Template3* and *Template4*) as controls, which contain only one promoter but otherwise identical elements including the *ynaJ-uspE* terminator as in *Template2* (**Figure 2A**). We observed that only 9% of restarted ECs on *Template3* released their RNAP and RNA at the terminator, while the majority fell within the *Retention* or *Readthrough* category (**Figure 2B-D**). On *Template4*, the *Release* events occurred at a higher frequency (39%) (**Figure 2E-G**), which is expected given the longer 3’ U tract of the terminator element in the reverse direction than in the forward direction (**Figure S3B**). Nevertheless, the termination efficiency was still much lower than that observed on *Template1* (**Figure 1G**), indicating that this terminator does not cause strong intrinsic termination in either direction.

**Figure 2.**
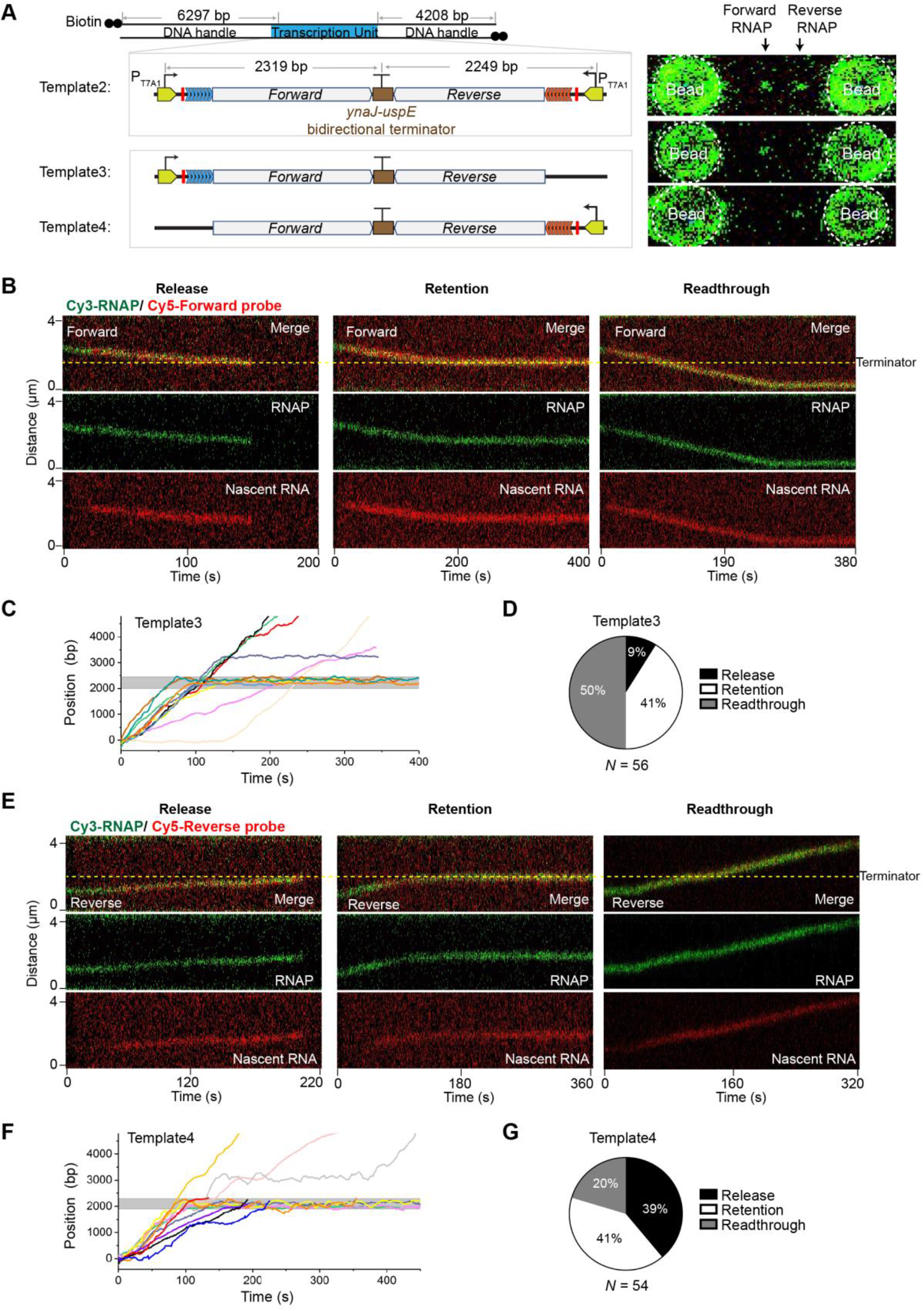
Unidirectional transcription on templates containing a bidirectional terminator. (**A**) (*Left*) Schematic of *Template2* that contains two opposing promoters and the *ynaJ*-*uspE* bidirectional terminator, *Template3* and *Template4* that contain the *ynaJ*-*uspE* bidirectional terminator but only one T7A1 promoter in the forward and reverse direction, respectively. (*Right*) 2D scan images showing Cy3-RNAPs at their respective stall sites on each template. (**B**) Representative kymographs showing a “Release” (*Left*), “Retention” (*Middle*), and “Readthrough” (*Right*) forward EC (RNAP in green, nascent RNA in red) at the bidirectional terminator on *Template3*. (**C**) Multiple overlaid trajectories of a forward RNAP translocating on *Template3* and displaying distinct behaviors at the terminator (gray zone). (**D**) Pie chart showing the fraction of forward EC release, retention, and readthrough events observed on *Template3*. (**E**) Representative kymographs showing a “Release” (*Left*), “Retention” (*Middle*), and “Readthrough” (*Right*) reverse EC (RNAP in green, nascent RNA in red) at the bidirectional terminator on *Template4*. (**F**) Multiple overlaid trajectories of a reverse RNAP translocating on *Template4* and displaying distinct behaviors at the terminator (gray zone). (**G**) Pie chart showing the fraction of reverse EC release, retention, and readthrough events observed on *Template4*. *N* denotes the total number of ECs. See also Figure S3.

Next, we examined the fate of ECs during convergent transcription using *Template2*. Satisfyingly, stalled ECs in both forward and reverse directions were observed to restart, translocate towards the terminator, and collide in a head-on manner within the termination zone (**Figure 3A** and **3B**). The collided RNAPs never bypassed each other, thus presenting a physical barrier against each other and efficiently preventing transcriptional readthrough, consistent with a previous atomic force microscopy study (Crampton et al., 2006). Indeed, the readthrough frequencies in both directions on *Template2* were significantly reduced compared to those observed on *Template3* and *Template4* that only support unidirectional transcription (**Figure 3C**). The occasional readthrough RNAP on *Template2* across the terminator was mostly blocked by an opposing RNAP at a distal site, preventing it from transcribing further (**Figure S4**).

**Figure 3.**
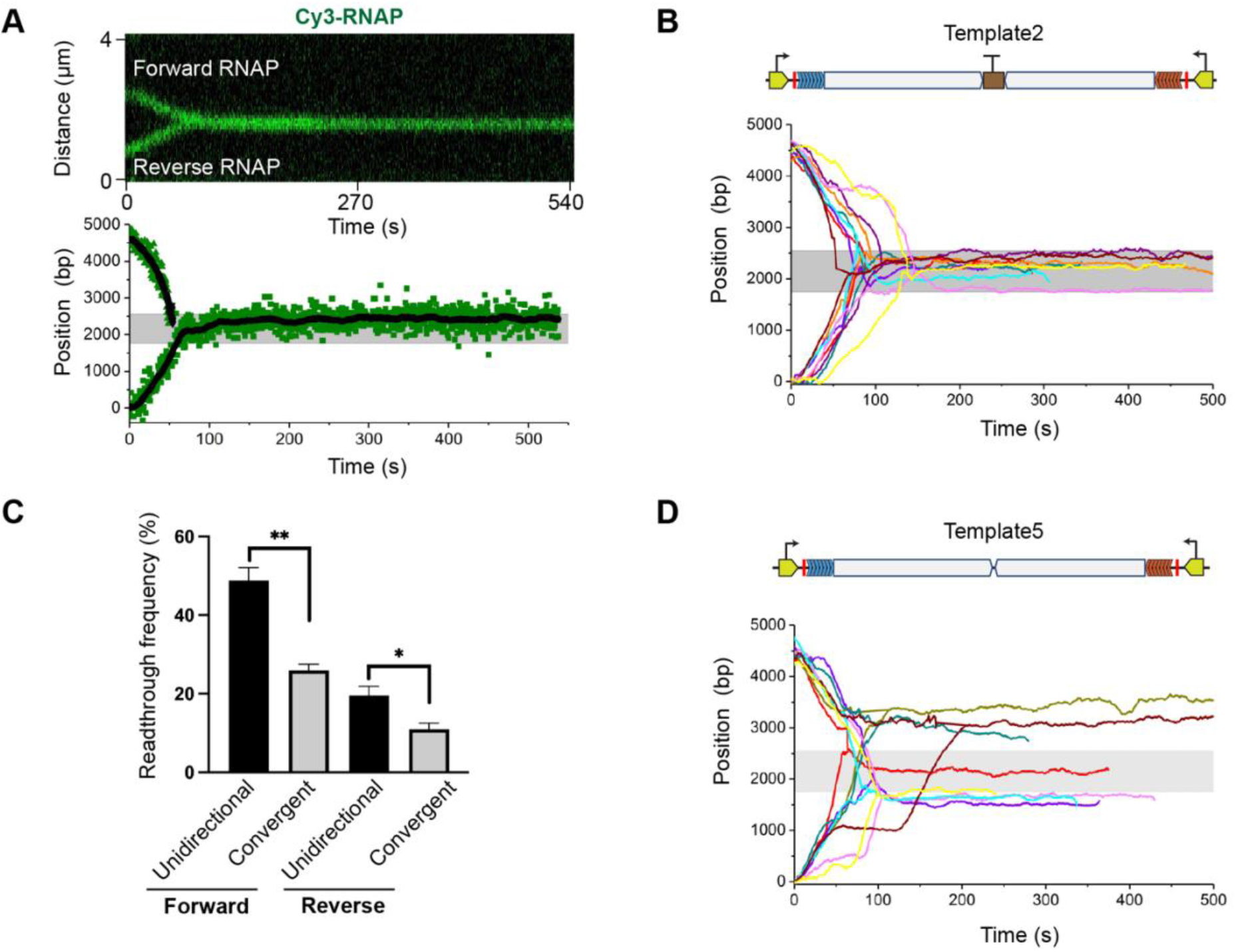
Direct visualization of convergent transcription. (**A**) (*Top*) A representative kymograph showing convergent transcription and head-on collision of two RNAPs. (*Bottom*) Extracted positions of the two RNAPs as a function of time for the same kymograph. Raw data and smoothed trajectories (± 10-s moving average of tracked points) are shown in green dots and black lines, respectively. Gray region indicates the bidirectional terminator position (2250 ± 400 bp). (**B**) Multiple overlaid trajectories of convergent RNAP pairs on *Template2* show collisions occur uniformly within the termination zone. (**C**) Frequency of RNAP reading through the *ynaJ*-*uspE* terminator in the forward or reverse direction observed on DNA templates that only allow unidirectional transcription (*Template3* and *Template4*; see Figure 2) versus on *Template2* that allows convergent transcription. Data are presented as mean ± SEM from three independent experiments. Significance was determined using two-sided unpaired Student’s *t*-tests (\**P* < 0.05; \*\**P* < 0.01). (**D**) Multiple overlaid trajectories of convergent RNAP pairs on *Template5* that lacks the bidirectional terminator show heterogeneous collision sites. See also Figures S4 and S5.

### Bidirectional terminator element is required for programmed collisions

How could the two convergent RNAPs initially located thousands of basepairs apart always collide at the same position? We postulated that the bidirectional terminator element—even though it does not mediate termination per se—may serve as a strong pausing signal for the converging ECs due to the RNA hairpin structure that it encodes. This would ensure that head-on collision events uniformly take place at the terminator site. The high *Retention* frequency observed on *Template3* and *Template4* supports this speculation (**Figure 2D** and **2G**). To further test it, we removed the terminator element from *Template2*, creating *Template5*. Interestingly, convergent transcription experiments using *Template5* yielded varied collision sites (**Figure 3D**). This result is corroborated by bulk in vitro transcription assays, which showed RNA products of well-defined sizes when the terminator element was present in the template but of heterogeneous lengths when the element was absent (**Figure S5**).

### Fate of transcription complexes after head-on collision

By monitoring the Cy3 fluorescence intensity, we noticed that in 56% of the head-on collision events, both RNAPs remained bound to the DNA after collision (i.e., Cy3 intensity doubled the pre-collision value) (**Figure 4A** *Left* and **Figure 4B**). Only a minor fraction of events showed one RNAP dissociated from the DNA (Cy3 intensity matched the pre-collision value) (39%; **Figure 4A** *Middle*) or both RNAPs dissociated after collision (Cy3 intensity dropped to zero) (5%; **Figure 4A** *Right*). We then performed three-color imaging to simultaneously track the fate of convergent RNAPs (by Cy3 fluorescence) and their respective nascent RNA (by AlexaFluor488 and Cy5 fluorescence). Again, in more than half of the collision events, the two converging ECs—including both RNAP and nascent RNA—were retained on the DNA (**Figure 4C** and **4D**). Therefore, even though the collided ECs effectively stop each other, the head-on collision itself is not sufficient to release the RNAP and nascent RNA. Other factors must be involved to complete the collision-driven termination process.

**Figure 4.**
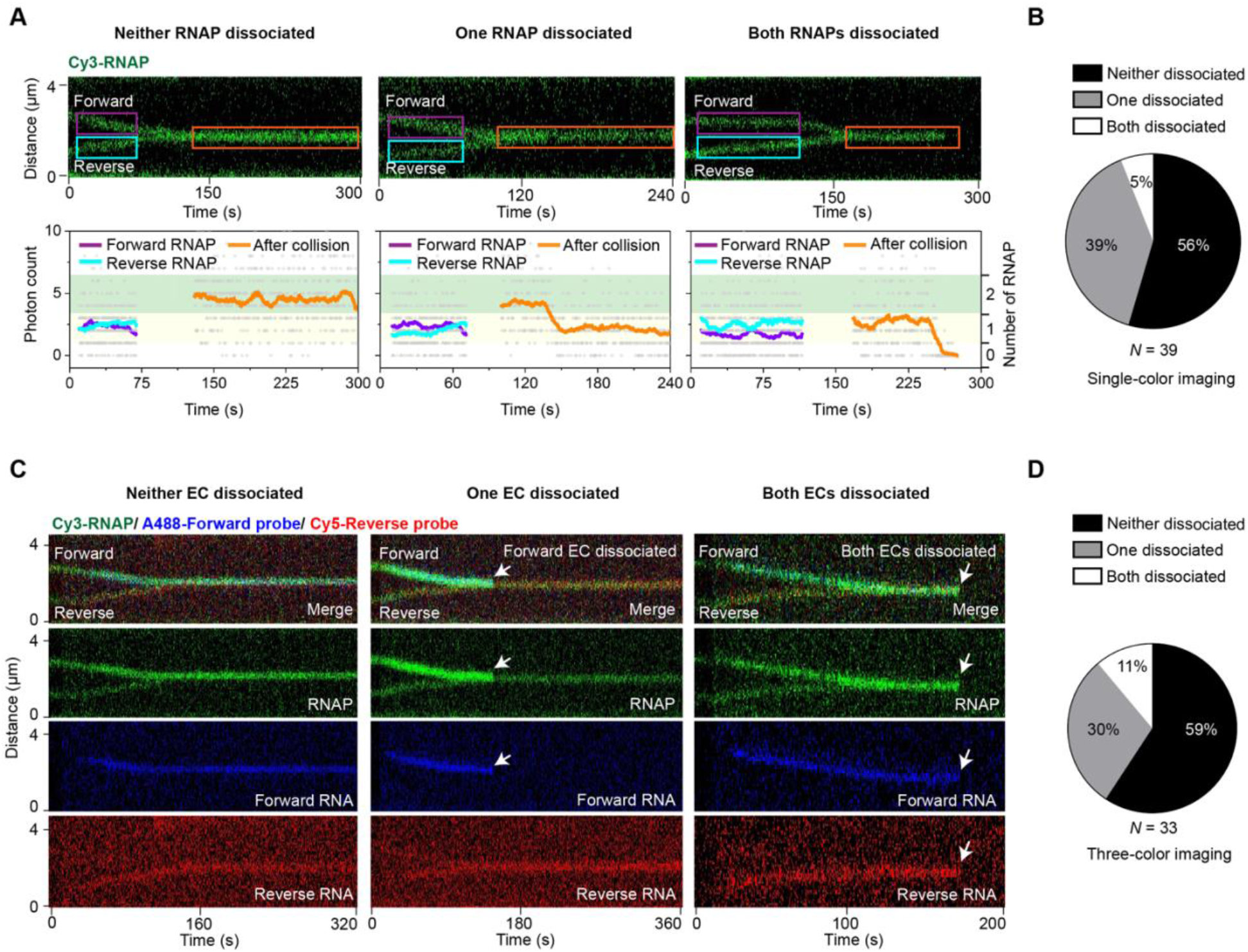
Outcome of head-on collisions at the bidirectional terminator. (**A**) Three different outcomes of head-on collision between two convergent RNAPs at the bidirectional terminator on *Template2* (depicted in Figure 2A). (*Top*) Representative kymographs showing both RNAPs remaining on DNA (*Left*), one of the RNAPs dissociated (*Middle*), or both RNAPs dissociated (*Right*) after collision. (*Bottom*) Cy3 intensity profiles for selected regions from the kymograph on top (colored boxes). The summed pixel intensities of individual frames were displayed as gray dots. The filtered values (± 10-s average of frames) were shown as colored lines corresponding to the colored boxes. The number of RNAPs within the selected regions were assigned based on the photon count: under 1 as zero RNAP; between 1 and 3.5 as one RNAP (yellow shade); between 3.5 and 6.5 as two RNAPs (green shade). (**B**) Pie chart showing the fraction of each outcome illustrated in (A). (**C**) Visualizing head-on collisions by three-color imaging (Cy3 for RNAP, AlexaFluor488 for forward RNA, Cy5 for reverse RNA). Representative kymographs showing both ECs remaining on DNA (*Left*), one of the ECs dissociated (*Middle*; forward EC dissociated in this example), or both ECs dissociated (*Right*) after collision. (**D**) Pie chart showing the fraction of each outcome illustrated in (C). *N* denotes the total number of collision events.

### Screening for factors that help accomplish termination

To identify such factors, we developed a bulk assay to evaluate the effect of several *E. coli* elongation and termination factors—namely Rho, NusA, NusG, Mfd, GreA and GreB (Washburn and Gottesman, 2015; Zenkin and Yuzenkova, 2015)—on the collision outcome. In this assay, transcription took place on biotinylated DNA templates immobilized on magnetic streptavidin beads (**Figure 5A**). Successfully released transcription complexes upon termination were washed away and the amount of retained complexes on the DNA was assessed by the intensity of fluorescent oligo probes on a gel. Consistent with our single-molecule results, we observed a much higher level of retained nascent RNA on the convergent transcription template (derived from *Template2*) than on the template with a strong intrinsic terminator (derived from *Template1*) (**Figure 5B**), again suggesting that head-on collisions alone cannot accomplish efficient termination. Of all the factors tested, only GreB induced a substantially reduced level of retained probe signal (**Figure 5C**). Notably, GreB reduced the retained probe signal only under the multi-round transcription condition; under single-round transcription condition (by adding rifampicin to prevent re-initiation), GreB exhibited little effect (**Figure 5D**). We then conducted single-molecule assays, which by design only enabled single-round transcription, and found that GreB did not change the collision outcome (i.e., most collided complexes still remained stable) (**Figure 5E**). GreB is a transcript cleavage factor known to rescue long backtracked RNAPs (Marr and Roberts, 2000; Shaevitz et al., 2003). Indeed, we found that GreB reduced the fraction of arrested ECs (**Figure S2**) in our single-molecule experiments (**Figure 5F**). Based on these results, we proposed that the role of GreB in collision-driven termination hinges not on its direct interaction with the collided RNAPs, but rather its ability to rescue arrested RNAPs and increase their “trafficking density” on DNA (**Figure S6**), thereby increasing the likelihood of a trailing RNAP running into a head-on collided complex.

**Figure 5.**
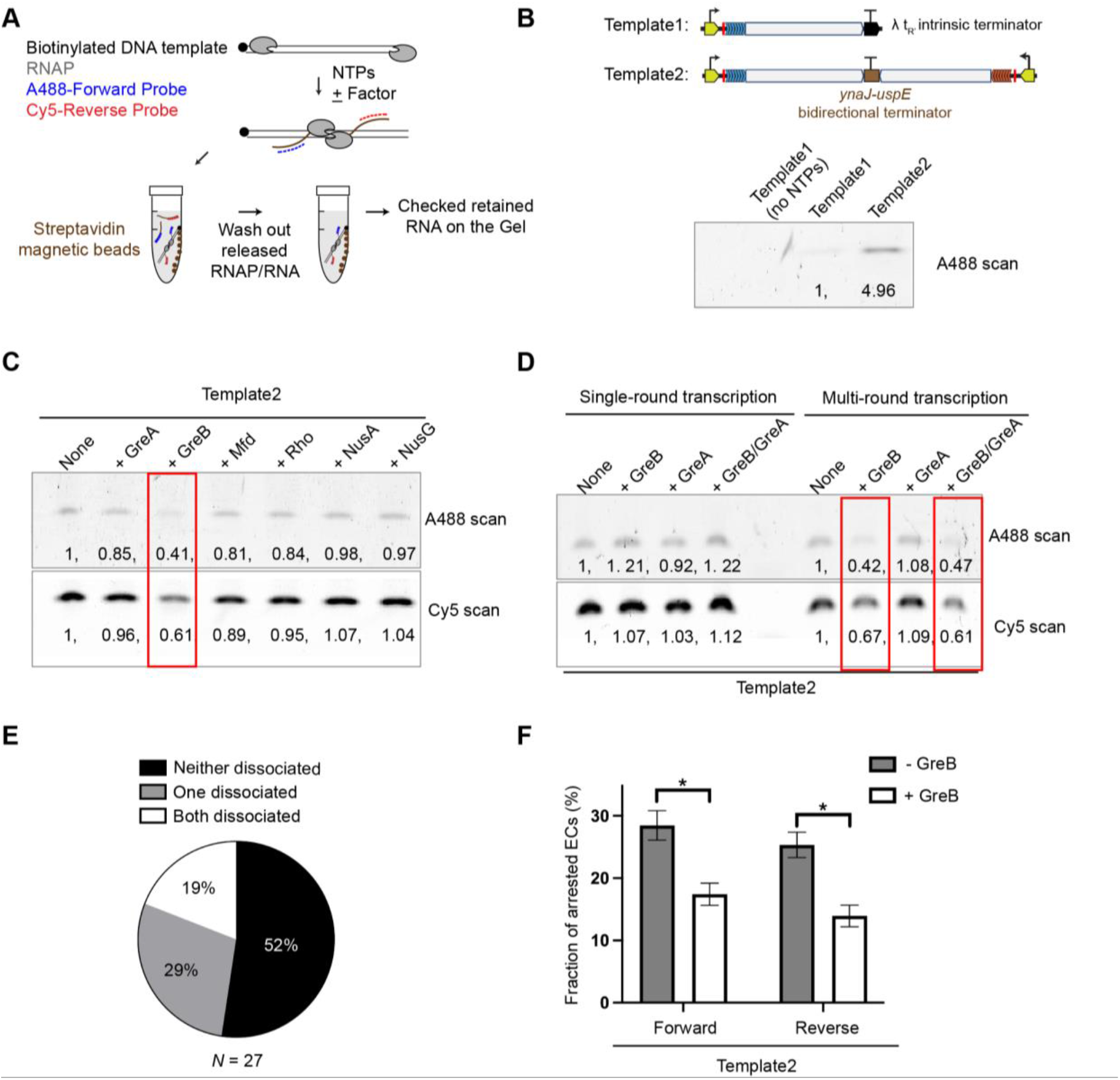
Screening for factors that facilitate the dissociation of head-on collided complexes. (**A**) Experimental workflow. Briefly, in vitro transcription reactions were performed on biotinylated DNA templates in the presence of fluorescently labeled probes to detect nascent RNA. After reaction, DNA substrates were pulled down by streptavidin magnetic beads. After thoroughly washing out the released RNAP and RNA products upon termination, the amount of RNA retained on the DNA template was assessed by the probe intensity from the gel. (**B**) Gel results for the assay described in (A) using DNA substrates derived from *Template1* (with an intrinsic terminator) and *Template2* (with a bidirectional terminator). AlexaFluor488-labeled forward probe intensities were normalized to the *Template1* value. (**C**) Gel results showing the effects of a panel of *E. coli* transcription factors on the amount of RNA products retained on *Template2*. AlexaFluor488-labeled forward probe and Cy5-labeled reverse probe intensities were normalized to the no-factor condition. The +GreB condition (outlined in red) exhibited reduced probe intensities. (**D**) Gel results showing the effect of GreB and/or GreA on the amount of retained RNA probes on *Template2* under the single-round or multi-round transcription condition. Probe intensities were normalized to the no-factor condition. (**E**) Pie chart showing the fraction of different outcomes for head-on collisions observed on *Template2* in the presence of GreB in the single-molecule assay. GreB was supplied in channel 4 at a final concentration of 800 nM. *N* denotes the total number of collision events. (**F**) Fraction of arrested forward and reverse ECs observed on *Template2* with or without GreB. Data are presented as mean ± SEM from three independent experiments. Significance was determined using two-sided unpaired Student’s *t*-tests (\**P* < 0.05). See also Figure S6.

### Multi-RNAP collision is key to bidirectional termination

To test the above proposed model further, we constructed a three-promoter template (*Template6*) by placing an additional T7A1 promoter upstream of the forward promoter in *Template2* (**Figure 6A**). This template allowed us to stall three ECs (two forward RNAPs, one reverse RNAP) and then restart them to observe both head-on and co-directional collisions in the single-molecule assay. Remarkably, most collisions at the bidirectional terminator that involved three ECs resulted in the release of two RNAPs from the DNA (i.e., the total Cy3 intensity after all three ECs collided was reduced to the single-RNAP level) (**Figure 6B**). Three-color experiments that simultaneously tracked the fate of convergent RNAPs and their respective nascent RNA also showed that the head-on collided complex—including both RNAPs and nascent RNAs—can be dislodged by an extra EC (**Figure 6C**). Overall, we observed that after both head-on and co-directional collisions occurred at the bidirectional terminator, the final number of RNAPs remaining on the DNA was predominantly one (67%), occasionally two (29%), rarely three (4%) (**Figure 6D**). These results strongly support our model that rear-ending by a trailing RNAP promotes dissociation of the head-on collided complex. In other words, heavy RNAP trafficking, counterintuitively, alleviates the traffic jam rather than exacerbating it. This model then predicts that the bidirectional termination efficiency would correlate positively with the RNAP “trafficking density” on DNA (i.e., more processive ECs in the presence of GreB or initiating from a stronger promoter). We thus analyzed our SEnd-seq dataset and indeed found that highly expressed gene pairs displayed significantly higher termination efficiencies than lowly expressed pairs (**Figure 6E** and **S7**).

**Figure 6.**
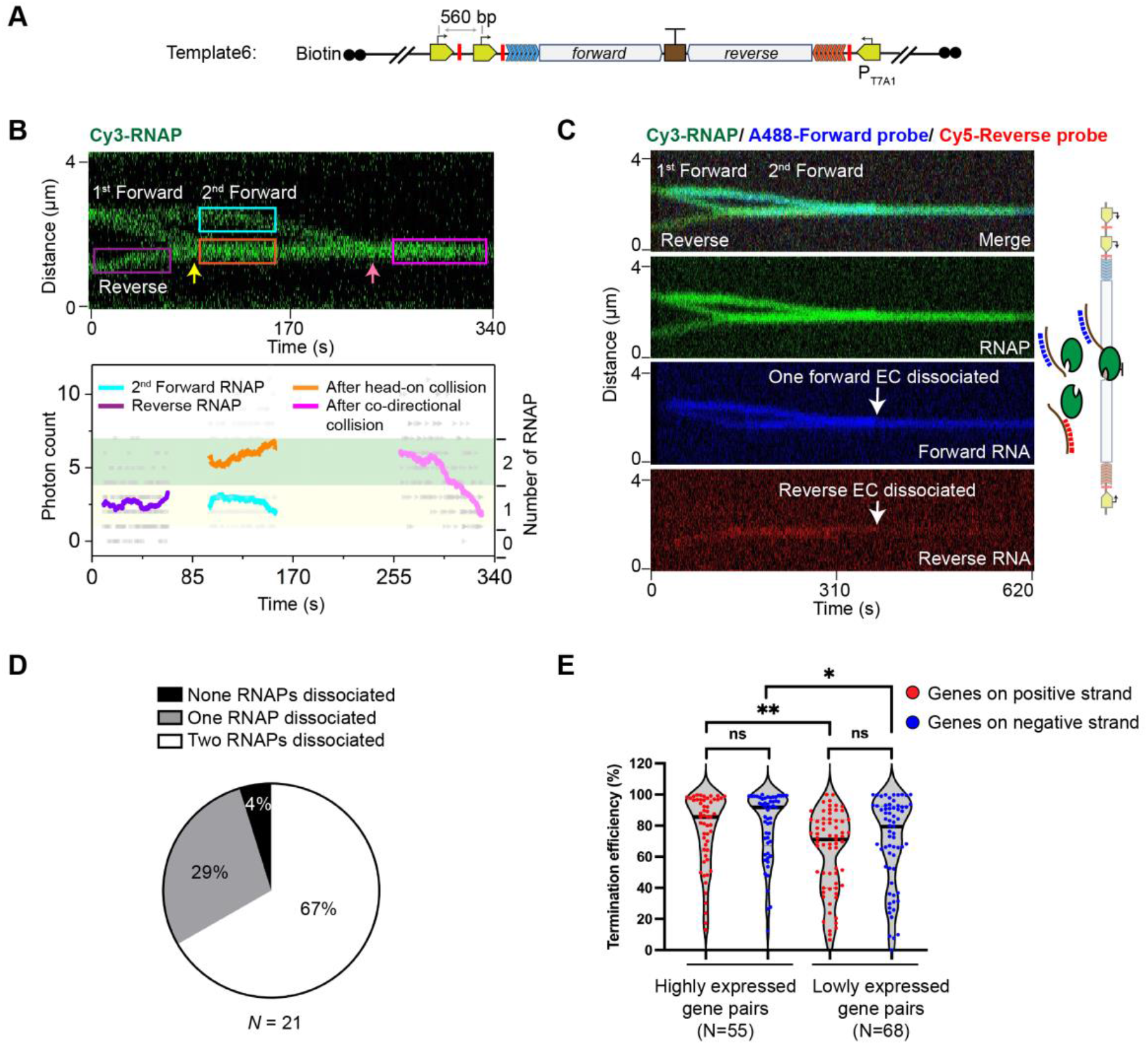
Outcome of sequential head-on and co-directional collisions at the bidirectional terminator. (**A**) Schematic of *Template6*, which is derived from *Template2* and contains an additional forward promoter and stall site. (**B**) A representative kymograph (*Top*) and corresponding Cy3 intensity profile (*Bottom*) showing head-on collision (yellow arrow) followed by co-directional collision (pink arrow) among two forward RNAPs and one reverse RNAP on *Template6*. The number of RNAPs for the selected regions (colored boxes) were analyzed and displayed in the same fashion as in Figure 4A. Two of the three RNAPs were eventually dissociated in this example. (**C**) A representative three-color kymograph (Cy3 for RNAP, AlexaFluor488 for forward RNA, Cy5 for reverse RNA) shows co-directional collision by the second forward EC dislodges both forward and reverse ECs in the head-on collided complex from *Template6* (AlexaFluro488 signal dropped by half and Cy5 signal disappeared at the white arrows). (**D**) Pie chart showing the fraction of different outcomes (none, one, or two of the three RNAPs dissociated) after sequential head-on and co-directional collisions at the bidirectional terminator on *Template6*. (**E**) Violin plot showing the distributions of bidirectional termination efficiencies for convergent *E. coli* gene pairs that are highly expressed (read counts > 100 for both genes) and lowly expressed (10 < read counts < 100 for both genes). *N* denotes the number of gene pairs in each group. Significance was determined using two-sided unpaired Student’s *t*-tests (ns, not significant; \**P* < 0.05; \*\**P* < 0.01). See also Figure S7.

## DISCUSSION

In this study, we described a model for bidirectional transcription termination that is collectively driven by head-on and co-directional RNAP collisions (**Figure 7**): 1) the bidirectional terminator element forms an RNA hairpin structure and thus traps the transcribing EC from either direction, which in turn serves as an efficient barrier to prevent the opposite EC from transcribing across the gene boundary. However, head-on collision itself is not sufficient to release the RNAPs and their associated RNA products; 2) a trailing RNAP subsequently runs into the head-on collided complex, releasing at least one of its two RNAPs and the nascent RNA; 3) the remaining RNAP continues to block any incoming opposite RNAP and the whole process repeats. This process occurs more frequently for highly expressed genes and is assisted by transcription factors like GreB. As such, this model suggests a direct link between the initiation efficiency and termination efficiency, adding to the functional roles of cooperative RNAP trafficking in transcriptional regulation (Epshtein and Nudler, 2003; Jin et al., 2010).

**Figure 7.**
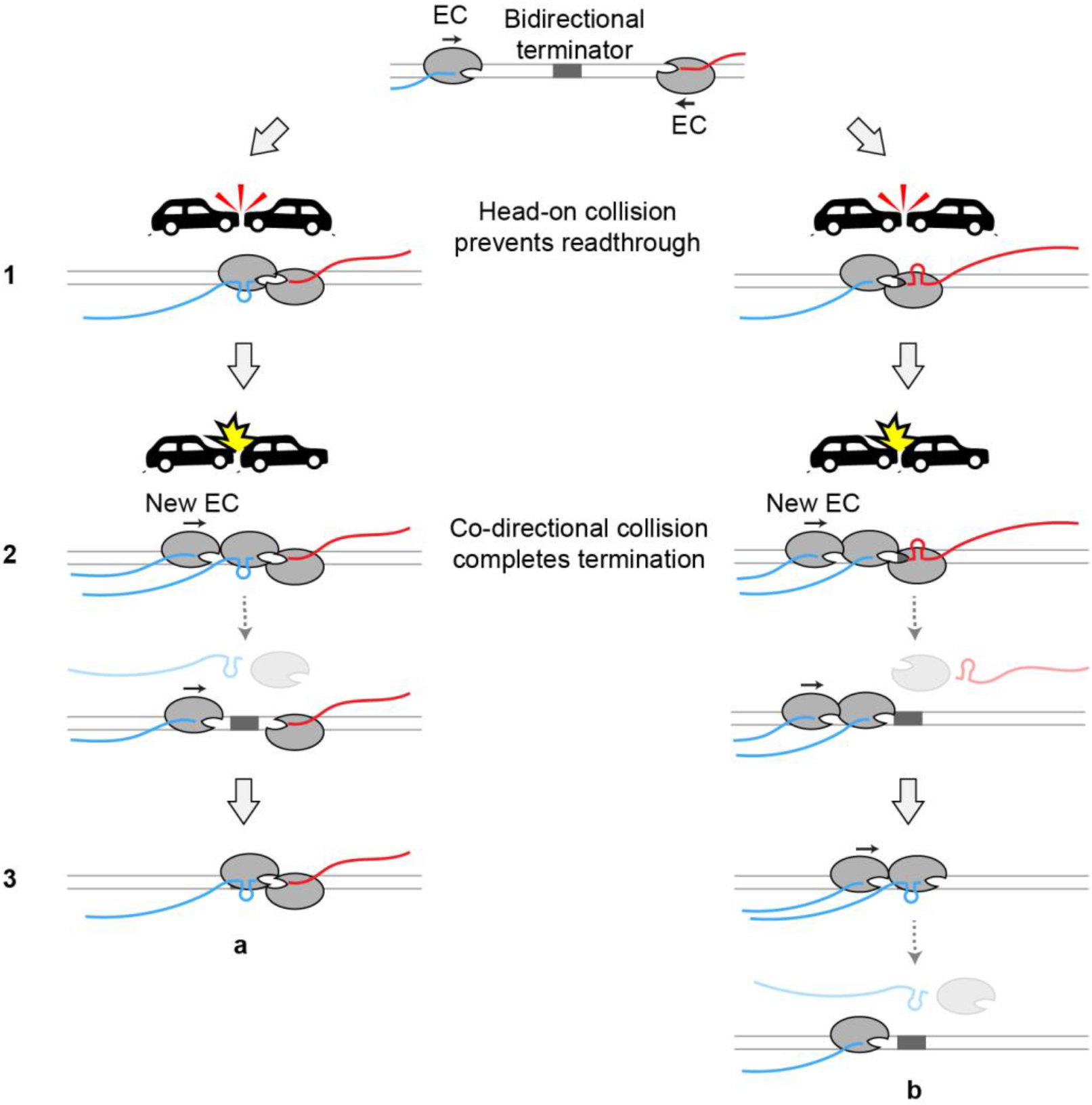
Proposed model for RNAP-collision-driven bidirectional transcription termination. The RNA hairpin structure formed at the overlapping region of the bidirectional terminator induces pausing of the EC from either direction. The opposite RNAP runs into the paused RNAP, forming a stable head-on collided complex that does not efficiently dissociate by itself (*Step 1*). When a trailing EC newly initiated from the promoter runs into the head-on collided complex, the RNAP that transcribed past the overlapping region is dislodged from the DNA along with its nascent RNA due to its lower stability induced by the RNA hairpin (*Step 2*). The trailing EC may either transcribes the overlapping region and forms a new head-on collided complex with the remaining EC (*Step 3*, *a*), or pushes its leading EC across the overlapping region and dislodges it (*Step 3*, *b*). Note that the new EC illustrated here travels in the forward direction. A reciprocal scenario where the new EC travels in the reverse direction is omitted for simplicity. This model explains the predominantly uniform 3’ ends of nascent transcripts observed at the bidirectional terminators (e.g., Figure S3A and S7).

Given the universal occurrence of convergent gene pairs and antisense transcripts in diverse organisms (Callen *et al.*, 2004; Georg and Hess, 2018; Gullerova and Proudfoot, 2012; Hobson *et al.*, 2012; Prescott and Proudfoot, 2002; Shearwin *et al.*, 2005; Yelin *et al.*, 2003), programmed RNAP collisions may represent an evolutionarily conserved mechanism for transcription termination. In *E. coli*, this mechanism is enabled by the strategic placement of bidirectional terminator elements, which is critical to shaping precise transcript 3’ boundaries. Future studies are required to assess the existence of similar pausing-prone elements between convergent genes in other organisms.

Collisions among RNAPs and among DNA-based motors were generally thought to be deleterious to genome stability. Our work suggests that motor collisions can in fact be harnessed to fine-tune gene expression. By enabling real-time observation of motor trafficking on DNA, our single-molecule platform will allow us to investigate a wide range of genomic conflicts and their potential physiological functions (Le and Wang, 2018).

### Limitation of the Study

Given the limited spatial resolution of our assay, we currently cannot determine whether RNAPs in the collided complex are in direct contact. We also did not explore how other processes, such as DNA supercoiling (Kim et al., 2019) and transcription-translation coupling (Proshkin et al., 2010), affect the outcome of collision-driven transcription termination. Future structural and single-molecule studies will provide insights into these questions.

## ACKNOWLEDGMENTS

We thank J. Chen, B. Malone, J. Brewer (Darst/Campbell Laboratory, Rockefeller University), R. Gong (Alushin Laboratory, Rockefeller University), and R. Mooney (Landick Laboratory, University of Wisconsin-Madison) for sharing reagents and technical assistance, and members of the Liu Laboratory for discussions. This work is supported by the Robertson Foundation, the Alfred P. Sloan Foundation Matter-to-Life Program, the Pershing Square Sohn Cancer Research Alliance, and an NIH Director’s New Innovator Award (DP2HG010510) to S.L.

## AUTHOR CONTRIBUTIONS

S.L. and L.W. conceived the project. L.W. performed experiments and data analysis. J.W.W. wrote scripts for single-molecule data analysis. X.J. provided and analyzed the SEnd-seq data. L.W. and S.L. wrote the manuscript with inputs from all authors.

## SUPPLEMENTAL INFORMATION

Supplemental Information includes seven figures and one table.

## STAR METHODS

### KEY RESOURCES TABLE

| REAGENT or RESOURCE | SOURCE | IDENTIFIER |
| --- | --- | --- |
| **Bacterial and Virus Strains** |  |  |
| *Escherichia coli* DH5α | Thermo Fisher | Cat# 18265017 |
| *Escherichia coli* BL21 (DE3) | Thermo Fisher | Cat# EC0114 |
| **Chemicals, Peptides, and Recombinant Proteins** |  |  |
| Alexa Fluor A488 NHS Ester | Thermo Fisher | Cat# A20000 |
| Cy3 maleimide mono-reactive dye | GE Healthcare | Cat# PA23031 |
| Cy5 NHS ester mono-reactive dye | GE Healthcare | Cat# PA15101 |
| RNaseOUT Recombinant ribonuclease inhibitor | Thermo Fisher | Cat# 10777019 |
| Gibson Assembly Master mix | New England Biolabs | Cat# E2611S |
| Klenow Fragment | New England Biolabs | Cat# M0212 |
| Biotin-11-dUTP | Thermo Fisher | Cat# R0081 |
| Biotin-14-dCTP | Thermo Fisher | Cat# 19518018 |
| Biotin-14-dATP | Thermo Fisher | Cat# 19524016 |
| Trolox | Sigma-Aldrich | Cat# 238813 |
| 4-Nitrobenzyl alcohol (NBA) | Sigma-Aldrich | Cat# N12821 |
| Cyclooctatetraene (COT) | Sigma-Aldrich | Cat# 138924 |
| Protocatechuate 3,4-Dioxygenase from *Pseudomonas* sp. (PCD) | Sigma-Aldrich | Cat# P8279 |
| 3,4-Dihydroxybenzoic acid (PCA) | Sigma-Aldrich | Cat# 37580 |
| Streptavidin Coated Polystyrene Particles (2.0-2.9 μm) | Spherotech | Cat# SVP-20-5 |
| Rifampicin | Sigma-Aldrich | Cat# R3501 |
| S6-RNAP | This paper | N/A |
| $\sigma^{70}$ | (Chen et al., 2019) | N/A |
| SFP synthase | (Zhou et al., 2007) | N/A |
| **Recombinant DNA** |  |  |
| pVS11-Sfp-rpoC (expressing S6-RNAP) | This paper | N/A |
| pLW83 (Template1) | This paper | N/A |
| pLW86 (Template2) | This paper | N/A |
| pLW87 (Template3) | This paper | N/A |
| pLW88 (Template4) | This paper | N/A |
| pLW89 (Template5) | This paper | N/A |
| pLW103 (Template6) | This paper | N/A |

| Software and Algorithms |  |  |
| --- | --- | --- |
| Bluelake | LUMICKS | https://lumicks.github.io/bluelake-api/ |
| Origin | OriginLab | https://www.originlab.com |
| PRISM | GraphPad Software | https://www.graphpad.com/scientific-software/prism/ |
| Python scripts for single-molecule data analysis | This paper | https://harbor.lumicks.com/ |

### RESOURCE AVAILABILITY

#### Lead contact

Requests for reagents and further information should be directed to the lead contact, Shixin Liu.

#### Materials availability

Plasmids generated in this study are available upon request to the lead contact.

#### Data and code availability

Scripts for single-molecule data analysis have been deposited to https://harbor.lumicks.com/ and are publicly free to download. Further information for raw data reported in this study is available from the lead contact upon request.

### EXPERIMENTAL MODEL AND SUBJECT DETAILS

#### Bacterial strains and growth conditions

The *Escherichia coli* DH5*α* was used for all cloning and plasmid extraction. The *E. coli* BL21 (DE3) was used for all protein expression. Cells were grown in LB medium containing 100 μg/mL ampicillin or 50 μg/mL kanamycin as necessary.

### METHOD DETAILS

#### DNA substrate preparation

##### DNA template for single-molecule experiments

The 2.2-kb-long transcription unit in *Template1* (**Table S1**) is adapted from pCDW114 (a gift from Jeff Gelles, addgene plasmid #70061), which was shown to allow processive transcription and robust detection of nascent RNA in a previous single-molecule study (Harden et al., 2016). It contains a λ P_R’_ promoter, a cassette harboring seven tandem repeats of a 21-bp sequence (5’-AGACACCACAGACCACACACA-3’), a portion of the *E. coli rpoB* gene, and a λ tR’ terminator. To make *Template1*, pCDW114 plasmid was digested with EcoRI/BamHI to remove its original λ P_R’_ promoter. A 171-bp DNA fragment that contains a stall site located at +39 position and flanking EcoRI/BamHI recognition sites (synthesized by GENEWIZ, Azenta Life Sciences) was then inserted between those restriction sites. Additionally, a 4-kb-long DNA fragment (PCR from *Myo7a* gene) and 5-kb-long DNA fragment (PCR from lambda genome) were inserted upstream and downstream, respectively, of the transcription unit by Gibson Assembly (New England BioLabs). The assembled 13.8 kb plasmid (pLW83) contains a unique XbaI endonuclease cutting site that generates a linear DNA upon XbaI digestion where the transcription unit is flanked between two asymmetric DNA handles (**Figure 1A**). The 5’ overhangs of the linearized plasmid were then filled in with biotinylated nucleotides by the 5’->3’ exonuclease-deficient DNA polymerase I Klenow fragment (New England BioLabs) to create terminally biotinylated DNA for optical trapping. The fill-in reaction was conducted by incubating 10 nM linearized plasmid DNA, 33 uM each of dGTP/biotin-14-dATP/biotin-11-dUTP/biotin-14-dCTP (Thermo Fisher), and 5 U Klenow in 1x NEB2 buffer at room temperature for 15 min. To stop the reaction, EDTA was added at a final concentration of 10 mM and the reaction mixture was heat inactivated at 75 °C for 20 min. The DNA was then ethanol precipitated overnight at −20 °C by 3x volumes of cold ethanol and 300 mM NaOAc pH 5.2. Precipitated DNA was recovered by centrifugation at 15,000 rpm at 4°C for 30 min. The DNA pellet was washed by 75% ethanol and then air-dried, resuspended in TE buffer (10 mM Tris-HCl pH 8.0, 1 mM EDTA) and stored at −20 °C.

To generate the *Template2* (**Figure 2A**), a 2.6-kb-long partial convergent transcription unit was synthesized by GENEWIZ (Azenta Life Sciences) into a pUC19 vector. The synthesized DNA contains the *ynaJ* gene, *ynaJ-uspE* overlapping termination sites, *uspE* gene and a nearby gene from *E. coli* genome, followed by the reverse repeat cassette (seven tandem repeats of a 19-bp sequence: 5’-CCCTATCCCTTATCTTAAC-3’) and a reverse T7A1 promoter with the same stall site as *Template1*. The whole fragment was then amplified and inserted into plasmid pLW83 to replace the λ tR’ intrinsic terminator by Gibson Assembly (New England BioLabs), yielding a full convergent transcription unit (**Table S1**). To better determine the orientation of the DNA substrate during experiments, the downstream DNA handle was truncated by replacing the 5-kb-long DNA fragment with 3.8-kb-long DNA fragment (PCR from lambda genome), resulting in a 15-kb plasmid (pLW86). Following XbaI digestion, a linear DNA containing the convergent transcription unit flanked by a 6.3-kb-long DNA handle and a 4.2-kb-long DNA handle on each side was produced (**Figure 2A**). Terminally biotinylated substrates were generated in the same way as described above.

To make the *Template3* and *Template4* (**Figure 2A**), either direction of the T7A1 promoter and repeat cassette were deleted from plasmid pLW86, yielding forward-only construct (plasmid pLW87) and reverse-only construct (plasmid pLW88). For *Template5* (**Figure 3D**), the *ynaJ-uspE* bidirectional terminator was deleted from plasmid pLW86, yielding a new plasmid (pLW89). For *Template6* (**Figure 6A**), an additional T7A1 promoter with a stall site was inserted into a location 560bp upstream of the forward T7A1 promoter of plasmid pLW86 by Gibson Assembly (New England BioLabs), yielding the three-promoter construct (plasmid pLW103). Linearized and terminally biotinylated DNA substrates were generated in the same way as described above.

##### DNA template for bulk transcription assay

The DNA templates used in **Figure S5B** were prepared from each corresponding construct by PCR with a universal pair of primers (LW8F: 5’-AACGCCAGCAACGCGGCCTTTTTACGGTTC /LW10R: 5’-CCTTATCTAAATTATAATGACGCACGATATG) that bind to the DNA handles region of the plasmid and amplify the transcription unit region. The biotinylated DNA templates used in **Figures 5B-D** were prepared by PCR with the same primer sequences (LW8F/LW10R) containing a 5’ biotin modification (IDT).

#### Protein and oligonucleotide probe labeling

To site-specifically label the *E. coli* RNAP core enzyme, we engineered a 12-residue S6 peptide tag (GDSLSWLLRLLN) (Zhou *et al.*, 2007) to the C-terminus of β’ subunit. Expression and purification of S6-RNAP (provided by Brandon Malone, Darst/Campbell Laboratory, Rockefeller University) followed the same procedure as previously described for wild-type RNAP (Chen *et al.*, 2019). For fluorescent labeling, the S6-RNAP protein, Sfp synthase and Cy3-CoA dye were incubated at a 1:2:5 molar ratio for 2 h at room temperature in the presence of 10 mM MgCl_2_. Excess dye and Sfp synthase were removed by a 100-kDa Amicon spin filter (Millipore). Final protein samples were aliquoted, flash frozen, and stored at −80 °C.

The oligonucleotide probes (forward: 5’-GTGTGTGGTCTGTGGTGTCT-3’; reverse: 5’-GTTAAGATAAGGGATAGGG-3’) were synthesized by IDT with an amino modification at their 5’ terminus. The oligos were mixed with AlexaFluor488 NHS ester (Thermo Fisher) or Cy5 NHS ester (Cytiva) at room temperature for 3 h. Excess free dye was subsequently removed by a Sephadex G-25 column (Cytiva).

#### Single-molecule experiments

##### Stalled elongation complex formed in tube

Cy3-labeled *E. coli* RNAP core enzyme was incubated with σ^70^ (provided by James Chen, Darst/Campbell Laboratory, Rockefeller University) at a 1:3 molar ratio for 2 h on ice. A 20-μl reaction mixture (4 μl of 100 ng/μl biotinylated DNA substrate, 5 μl of 3 μM Cy3-RNAP-σ^70^ holoenzyme, 4 μl of 5x Transcription buffer, 0.5 μl of RNaseOUT Inhibitor and nuclease-free water) was incubated at 37 °C for 10 min. Transcription was initiated by the addition of 50 μM of ApU dinucleotide (TriLink), 150 μM of ATP, UTP and GTP (New England Biolabs) and subsequent incubation for 10 min at 37 °C. Heparin (Sigma-Aldrich) was then added to a final concentration of 100 μg/ml for another 5-min incubation at 37 °C. The final reaction mixture was diluted with 500 μl 1x transcription buffer (25 mM Tris-HCl pH 7.5, 150 mM KCl, 10 mM MgCl_2_, 1mM DTT) to load into the instrument.

##### Single-molecule data collection

Single-molecule experiments were performed at room temperature on a LUMICKS C-Trap instrument. Channels of the microfluidics chip were first passivated by flowing BSA (0.1% w/v in PBS, Sigma-Aldrich) and then Pluronic F127 (0.5% w/v in PBS, Sigma-Aldrich) into the flow cell for 10 min each. Streptavidin-coated polystyrene beads (2.07-μm diameter, Spherotech), pre-stalled ECs on the biotinylated DNA substrate, imaging buffer, imaging buffer with all the four ribonucleotides (1 mM of each) were loaded into laminar-flow-separated channels 1-4, respectively. Imaging buffer included an oxygen scavenging system [10 nM protocatechuate-3,4-dioxygenase (Sigma-Aldrich) and 2.5 mM protocatechuic acid (Sigma-Aldrich)], as well as a triple-state quenching cocktail [1 mM cyclooctatetraene (Sigma-Aldrich), 1 mM 4-nitrobenzyl alcohol (Sigma-Aldrich) and 1 mM Trolox (Sigma-Aldrich)] in transcription buffer. A single DNA tether was caught between the streptavidin beads in the optical traps, and the tether was held under 5 pN of tension. The tether was then moved to channel 3 for confocal scanning using the 532 nm laser to verify the existence of stalled EC. Naked DNA or protein aggregates were discarded during this step. A DNA tether containing the stalled EC was then moved to channel 4 to resume transcription. When needed, AlexaFluor488 or Cy5-labeled forward probe and/or reverse probe (5 nM of each) were included in channel 4 to detect nascent RNA as specified in the text. Kymographs were generated via confocal line scanning through the center of the two beads at 0.32 s/line (pixel time: 0.1 ms). Data acquisition was carried out using Bluelake, the commercial software included with the LUMICKS C-trap.

#### Single-molecule data analysis

Single-molecule force and fluorescence data from the.h5 files generated from Bluelake were analyzed using tools in the lumicks.pylake Python library supplemented with other Python modules in a custom GUI Python script titled “C-Trap.h5 File Visualization GUI” (http://harbor.lumicks.com/single-script/c5b103a4-0804-4b06-95d3-20a08d65768f), which was used to export the confocal scans or kymographs in TIFF format, as well as to extract the real-time force, distance and confocal photon counts.

##### Extracting the positions of translocating RNAP

Positions of Cy3-RNAP along the DNA template as a function of time were extracted from kymographs by a home-written Python script titled “kymotracker_calling_script.py” (https://harbor.lumicks.com/single-script/4db9d63e-1f93-0c0-9b90-e99066469578), which applied the lumicks.pylake Python package’s greedy line tracking algorithms to define line traces. In general, a pixel threshold of 1 photon count, line width of 5 pixels, window of 20 frames, minimum line length of 50 frames provided the best tracking of moving RNAPs, but these parameters were slightly altered on a trace-by-trace basis to optimize the line tracking. Extracted trajectories were further smoothed by averaging adjacent ± 10 s time points of each position. Individual trajectories were then aligned based on the starting position of halted RNA polymerases. Traveling distance (nm) was converted to base-pair using the force-extension characteristics of the DNA substrate (0.327 nm/bp under 5pN tension). For the convergent transcription substrate (*Template2*), detection errors were inevitable during line tracking when two converging RNAPs come in close proximity due to the overlap of their fluorescent signal, resulting in discontinued lines.

##### Extracting the intensity of translocating RNAP

A custom Python script titled “area_photon_count_extractor.py” (https://harbor.lumicks.com/single-script/23f33367-7b02-4762-9457-37348da59194) was used to extract intensity of the lines at user-defined regions to compare the intensity before and after collision. This method of intensity analysis provided more reliable results than summing the photon counts around the tracked lines because the line tracking analysis biases selection of frames by setting a pixel threshold. The summed photon counts across each line scan within the selected regions were smoothed by averaging adjacent ± 10 s time points of the individual scan values and plotted in OriginLab.

#### Bulk transcription assay

The general reaction mixture contained 0.5 μg DNA template, 0.5 μl *E. coli* RNAP holoenzyme (New England Biolabs), 4 μl of 5x reaction buffer (New England Biolabs), 0.5 μl of RNaseOUT Inhibitor (Thermo Fisher) and nuclease-free water to a final volume of 20 μl.

##### Detecting RNA products

The mixture was incubated at 37 °C for 10 min and then 1 μl of 20 mM ribonucleotide mix (New England Biolabs) was added to initiate transcription. After 10 min of incubation at 37 °C, the DNA template and protein were digested with 0.3 μl of TURBO DNase (Thermo Fisher) for 10 min and then 0.5 μl of Protease K (Thermo Fisher) for 10 min subsequently. Final reactions were supplied with 2x RNA loading buffer (95% Formamide, 0.02% SDS, 1 mM EDTA) and separated by 1% agarose gel in 0.5x TBE buffer. The gel was stained by SYBR Gold (Thermo Fisher) and scanned by Axygen Gel System (Corning).

##### Probing the effect of factors on releasing head-on collided ECs

The expression plasmids for Rho, NusA, NusG, GreA and GreB were provided by Rachel Anne Mooney (Landick Laboratory, University of Wisconsin, Madison), and the proteins were purified by Rui Gong (Alushin Laboratory, Rockefeller University) following the protocols described previously (Hao et al., 2021; Perederina et al., 2006). The Mfd protein was provided by Joshua Brewer (Darst/Campbell Laboratory, Rockefeller University). After incubating the reaction mixture at 37 °C for 10 min, indicated factors were added at a final concentration of 600 nM and incubated for another 5 min at 37 °C. Then 1 μl of 20 mM ribonucleotide mix, 0.5 μl of 2 μM AlexaFluor488-forward probe and 0.5 μl of 2 μM Cy5-reverse probe were added to the reaction and incubated for 10 min at 37 °C. After transcription, 20 μl of streptavidin magnetic beads (New England Biolabs) were added to the reaction and mixed on a rotator for 20 min at room temperature. Released RNA and RNAP were washed out by 300 μl of 1x transcription buffer (25 mM Tris-HCl pH 7.5, 150 mM KCl, 10 mM MgCl_2_, 1mM DTT) three times. Final reactions were supplied with 2x RNA loading buffer and loaded to 8% Urea-PAGE after boiling at 90 °C for 5 min. The gel was scanned by Typhoon FLA 7000 (GE Healthcare) and quantified by ImageJ.

##### Single-round transcription

The mixture was incubated at 37 °C for 10 min and then stalled ECs were formed by adding 50 μM of ApU dinucleotide (TriLink), 150 μM of ATP, UTP and GTP (New England Biolabs) for another 10 min at 37 °C. Transcription was continued by supplying 1 mM ribonucleotide mix and 50 μg/ml rifampicin (Sigma Aldrich) which blocks re-initiation. The reaction was incubated at 37 °C for 10 min in the presence of fluorescently labeled probes and the following procedures were the same as above.

#### SEnd-seq data analysis

Overlapping bidirectional termination sites were identified by SEnd-seq as described previously (Ju *et al.*, 2019). Highly expressed convergent gene pairs were defined as those with >100 read counts for each gene in the pair; lowly expressed gene pairs were those with 10 < read counts <100 for each gene in the pair. Termination efficiency for both positive and negative directions of these gene pairs were plotted in GraphPad Prism.

### QUANTIFICATION AND STATISTICAL ANALYSIS

The number of molecules or events analyzed is indicated in the text or figure legends. Errors reported in this study represent the standard error of the mean unless noted otherwise. *P* values were determined using the two-sided, unpaired, Student’s *t*-tests with GraphPad Prism. The difference between two groups was considered statistically significant when *P* < 0.05 (\**P* < 0.05; \*\**P* < 0.01; \*\*\**P* < 0.001; ns, not significant).

