## Supplementary for "Head-on and co-directional RNA polymerase collisions orchestrate bidirectional transcription termination"

### SUPPLEMENTAL FIGURES AND TABLES

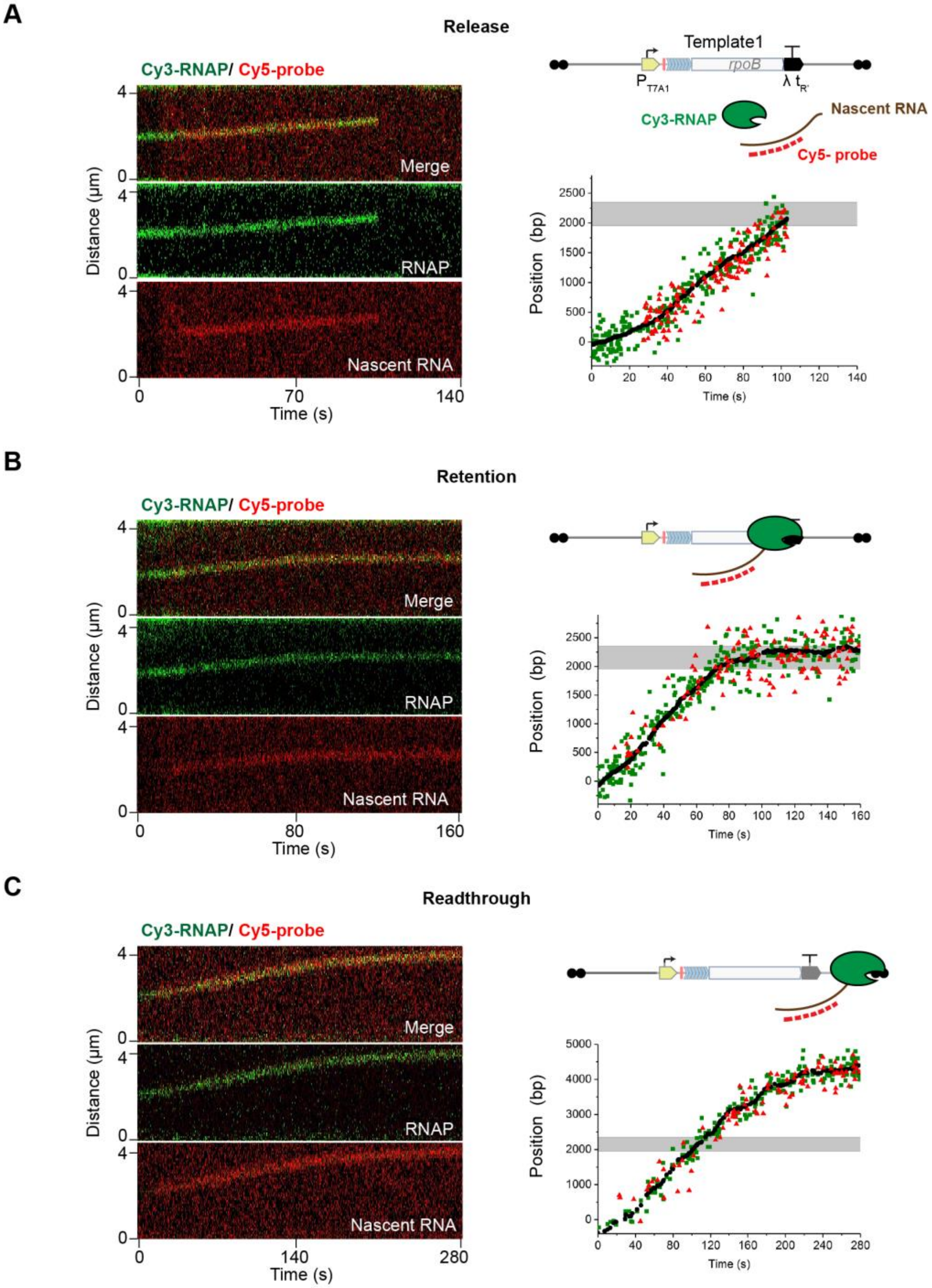

**Figure S1. Fate of the transcription complex at an intrinsic terminator, Related to Figure 1** (**A**) A "Release" example. (*Left*) A representative kymograph showing immediate release of RNAP (green) and nascent RNA (red) when the EC arrived at the λ tR' intrinsic terminator on *Template1* (see Figure 1A). (*Right*) Positions of RNAP (green) and nascent RNA (red) on the DNA template as a function of time extracted from the left kymograph.

(**B**) A “Retention” example. A representative kymograph (*Left*) and extracted RNAP/nascent RNA positions (*Right*) showing retention of the EC at the terminator.
(**C**) A “Readthrough” example. A representative kymograph (*Left*) and extracted RNAP/nascent RNA positions (*Right*) showing readthrough of the EC across the terminator. Raw data and smoothed RNAP trajectories (± 10-s moving average of tracked points) are shown in scattered dots and black lines, respectively. Gray regions indicate the terminator position.

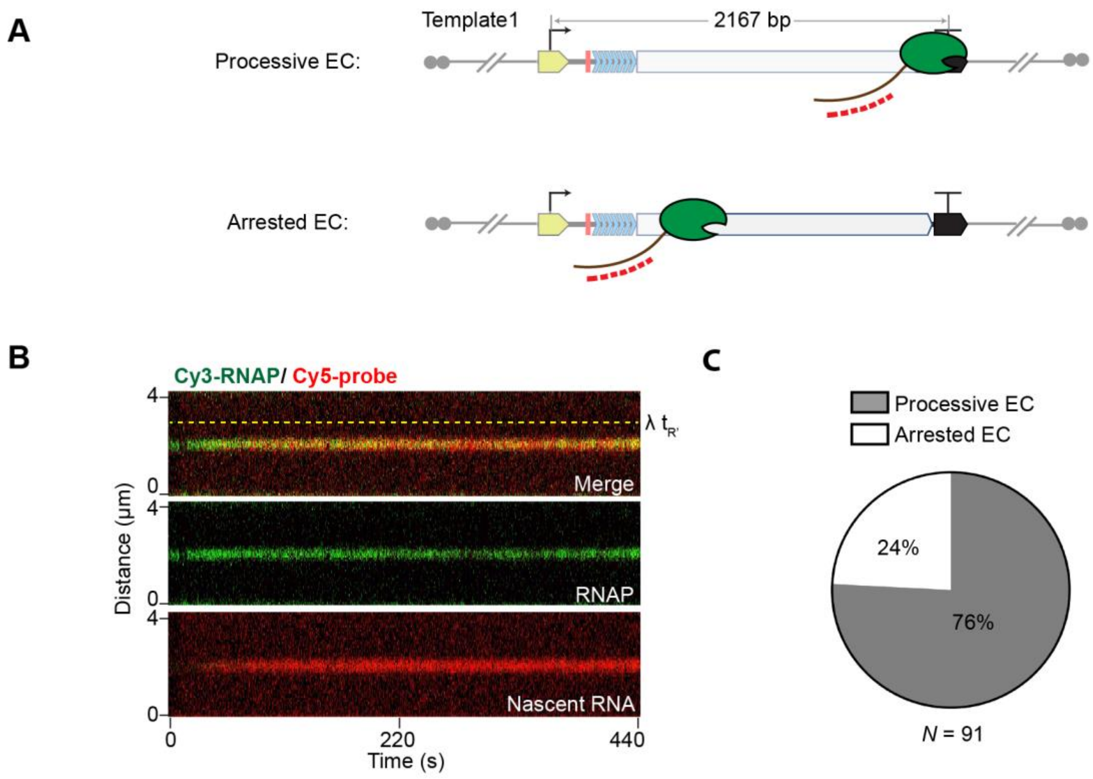

**Figure S2. Arrested elongation complexes, Related to Figure 1**

(**A**) Schematics of a processive EC (*Top*) that successfully reached the terminator site and an arrested EC (*Bottom*) that failed to reach the terminator.

(**B**) A representative two-color kymograph for arrested ECs showing halted translocation of the RNAP (green) and nascent RNA (red) before reaching the terminator on *Template1*.

(**C**) Pie chart showing the fraction of processive versus arrested ECs observed on *Template1*. *N* denotes the total number of ECs.

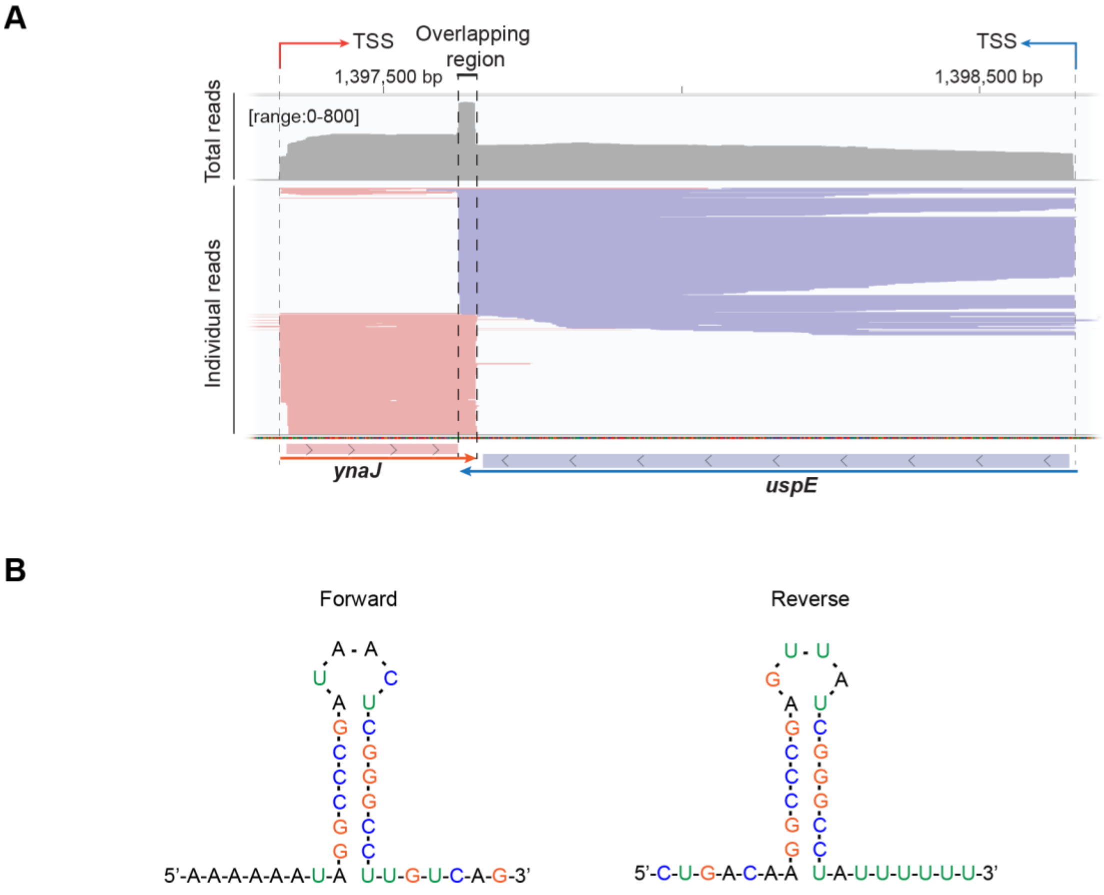

**Figure S3. Bidirectional terminator located between *ynaJ* and *uspE* genes in the *E. coli* genome, Related to Figure 2**

(**A**) SEnd-seq data track for the convergent *ynaJ*-*uspE* gene pair exhibiting a bidirectional terminator, which was incorporated in *Template2*, *Template3*, *Template4*, and *Template6*.

(**B**) Predicted secondary structure for the RNA transcribed from the overlapping region of the *ynaJ*-*uspE* bidirectional terminator in both forward and reverse directions.

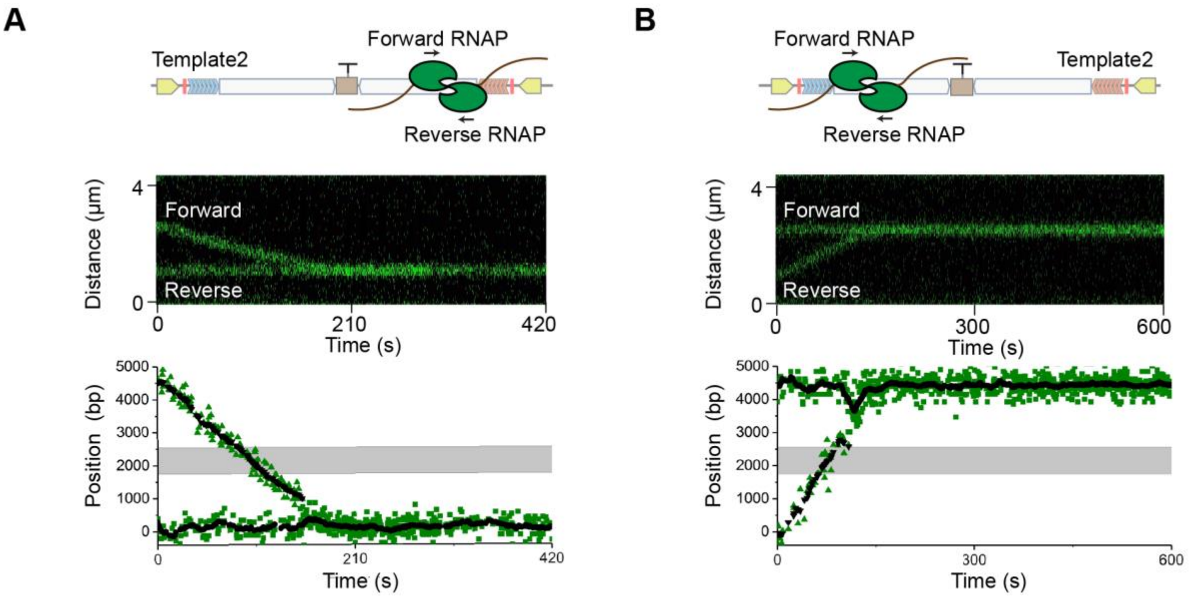

**Figure S4. Readthrough collisions, Related to Figure 3**

(**A**) An example in which a forward RNAP translocated across the *ynaJ-uspE* bidirectional terminator on *Template2* (see Figure 2A) and collided into an arrested reverse RNAP near the reverse stall site.

(**B**) An example in which a reverse RNAP translocated across the terminator and collided into an arrested forward RNAP near the forward stall site. *Top*: schematics; *Middle*: kymographs; *Bottom*: extracted RNAP positions on the template (green dots represent raw data; black dots represent ± 10-s moving averages of raw data points; and gray shades represent the termination zone).

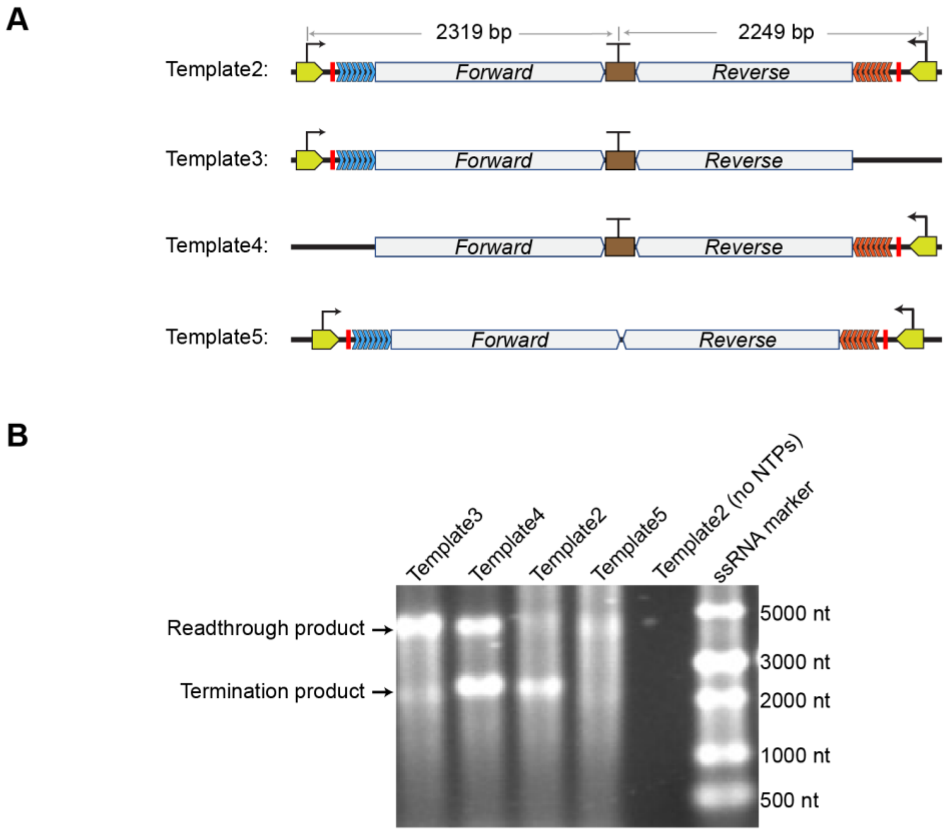

**Figure S5. Bulk transcription assays for evaluating termination and readthrough RNA products, Related to Figure 3**

(**A**) Schematics of DNA templates for unidirectional transcription (*Template3* and *Template4*) and convergent transcription (*Template2* and *Template5*). *Template5* lacks the bidirectional terminator element.

(**B**) Agarose gel results showing significant transcriptional readthrough on *Template3* and *Template4*, reduced readthrough and uniform termination on *Template2*, and varied RNA product lengths from *Template5*.

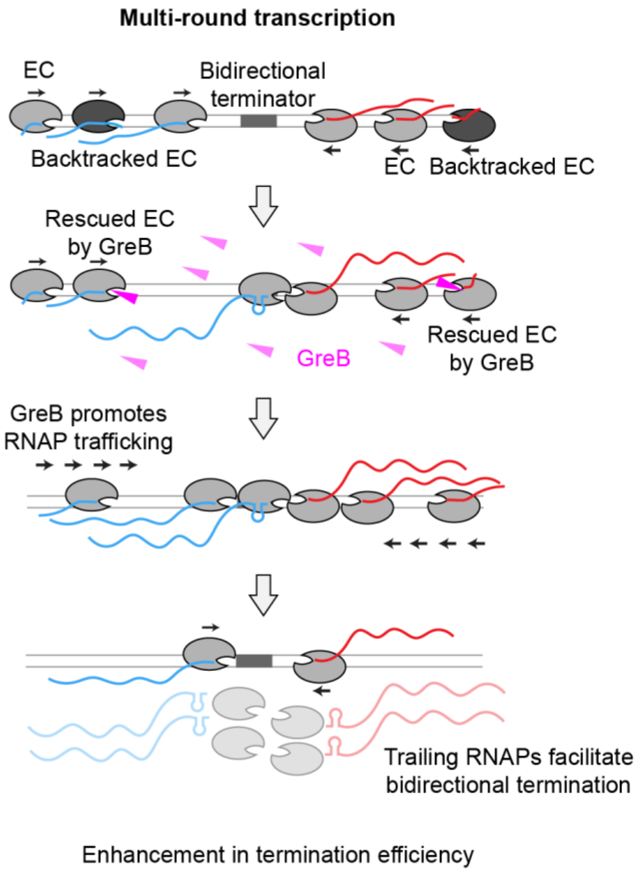

**Figure S6. Proposed model for GreB's effect on head-on collided ECs, Related to Figure 5**
GreB rescues long-backtracked RNAPs and enhances the RNAP trafficking density on DNA. When multi-round transcription is allowed, there is a higher chance of a trailing RNAP running into a head-on collided complex in the presence of GreB than in its absence, which facilitates dissociation of the collided complex. In the case of single-round transcription, this effect is not observed since there is no trailing RNAP and GreB does not directly destabilize the collided complex.

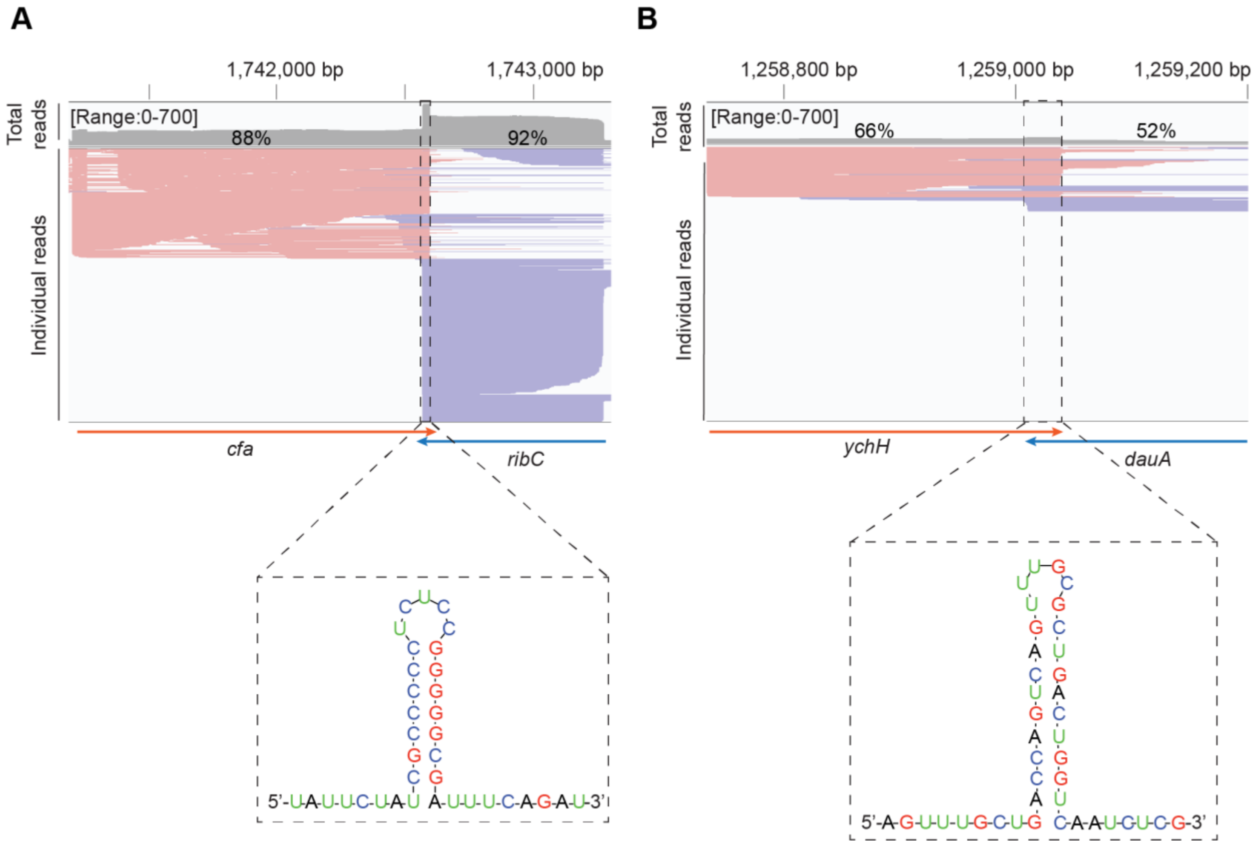

**Figure S7. Correlation between the expression level and termination efficiency of convergent gene pairs, Related to Figure 6**

(**A**) SEnd-seq data track for a highly expressed convergent gene pair (*cfa-ribC*) showing high termination efficiencies (88% for *cfa* and 92% for *ribC*).

(**B**) SEnd-seq data track for a lowly expressed convergent gene pair (*ychH-dauA*) showing lower termination efficiencies (66% for *ychH* and 52% for *dauA*). Predicted secondary structures for the RNA transcribed from the overlapping region of the bidirectional terminator (on the positive strand) are shown underneath the SEnd-seq reads.

**Table S1. DNA template sequences, Related to Figures 1—6**

| Name | Sequence | Notes |
|---|---|---|
| *Template1* | AAGATTAATTTAAAATTTATCAAAAAGAGTATTGACTTAA<br>AGTCTAACCTATAGGATACTTACAGCCATATGAGAGTTG<br>AAGTGGAATGGAGAGAATGAAGGATACCTAGGATCCagac<br>accacagaccacacacaagacaccacagaccacacacaagacaccacagaccacacacaa<br>gacaccacagaccacacacaagacaccacagaccacacacaagacaccacagaccacaca<br>caagacaccacagaccacacacagcatgcaacctgttcgtacgtatcgaccgtcgccgtaaac<br>tgcctgcgaccatcattctgcgcgccctgaactacaccacagagcagatcctcgacctgttcttt<br>gaaaaagttatctttgaaatccgtgataacaagctgcagatggaactggtgccggaacgcctgc<br>gtggtgaaaccgcatcttttgacatcgaagctaacggtaaagtgtacgtagaaaaaggccgcc<br>gtatcactgcgcgccacattcgccagctggaaaaagacgacgtcaaactgatcgaagtcccgg<br>ttgagtacatcgcaggtaaagtggttgctaaagactatattgatgagtctaccggcgagctgatct<br>gcgcagcgaacatggagctgagcctggatctgctggctaagctgagccagtctggtcacaag<br>cgtatcgaaacgctgttcaccaacgatctggatcacggcccatatatctctgaaaccttacgtgtc<br>gacccaactaacgaccgtctgagcgcactggtagaaatctaccgcatgatgcgccctggcga<br>gccgccgactcgtgaagcagctgaaagcctgttcgagaacctgttcttctccgaagaccgttat<br>gacttgtctgcggttggtcgtatgaagttcaaccgttctctgctgcgcgaagaaatcgaaggttcc<br>ggtatcctgagcaaagacgacatcattgatgttatgaaaaagctcatcgatatccgtaacggtaa<br>aggcgaagtcgatgatatcgaccacctcggcaaccgtcgtatccgttccgttggcgaaatggc<br>ggaaaaccagttccgcgttggcctggtacgtgtagagcgtgcggtgaaagagcgtctgtctctg<br>ggcgatctggataccctgatgccacaggatatgatcaacgccaagccgatttccgcagcagtg<br>aaagagttcttcggttccagccagctgtctcagtttatggaccagaacaacccgctgtctgagatt<br>acgcacaaacgtcgtatctccgcactcggcccaggcggtctgacccgtgaacgtgcaggcttc<br>gaagttcgagacgtacacccgactcactacggtcgcgtatgtccaatcgaaacccctgaaggt<br>ccgaacatcggtctgatcaactctctgtccgtgtacgcacagactaacgaatacggcttccttga<br>gactccgtatcgtaaagtgaccgacggtgttgtaactgacgaaattcactacctgtctgctatcga<br>agaaggcaactacgttatcgcccaggcgaactccaacttggatgaagaaggccacttcgtaga<br>agacctggtaacttgccgtagcaaaggcgaatccagcttgttcagccgcgaccaggttgactac<br>atggacgtatccacccagcaggtggtatccgtcggtgcgtccctgatcccgttcctggaacacg<br>atgacgccaaccgtgcattgatgggtgcgaacatgcaacgtcaggccgttccgactctgcgcg<br>ctgataagccgctggttggtactggtatggaacgtgctgttgccgttgactccggtgtaactgcg<br>gtagctaaacgtggtggtgtcgttcagtacgtggatgcttcccgtatcgttatcaaagttaacgaa<br>gacgagatgtatccgggtgaagcaggtatcgacatctacaacctgaccaaatacacccgttcta<br>accagaacacctgtatcaaccagatgccgtgtgtgtctctgggtgaaccggttgaacgtggcga<br>cgtgctggcagacggtccgtccaccgacctcggtgaactggcgcttggtcagaacatcgCT<br>TACTACCGATTCCGCCTAGTTGGTCACTTCGACGTATCGT<br>CTGGAACTCCAACCATCGCAGGCAGAGAGGTCTGCAAAA<br>TGCAATCCCGAAACAGTTCGCAGGTAATAGTTAGAGCCT<br>GCATAACGGTTTCGGGATTTTTTATATCTGCACAACAGGT<br>AAGAGCATTGAGTCGATAATCG | Underline: T7A1 promoter<br><br>Red: stall site<br><br>Blue: 7 tandem repeats (*agacaccacag accacacaca*)<br><br>Gray: 1.8-kb transcribed region<br><br>Brown: λ tR' intrinsic terminator |
| | AAGATTAATTTAAAATTTATCAAAAAGAGTATTGACTTAA<br>AGTCTAACCTATAGGATACTTACAGCCATATGAGAGTTG<br>AAGTGGAATGGAGAGAATGAAGGATACCTAGGATCCagac<br>accacagaccacacacaagacaccacagaccacacacaagacaccacagaccacacacaa<br>gacaccacagaccacacacaagacaccacagaccacacacaagacaccacagaccacaca<br>caagacaccacagaccacacacagcatgcaacctgttcgtacgtatcgaccgtcgccgtaaac<br>tgcctgcgaccatcattctgcgcgccctgaactacaccacagagcagatcctcgacctgttcttt<br>gaaaaagttatctttgaaatccgtgataacaagctgcagatggaactggtgccggaacgcctgc | Underline: forward T7A1 promoter<br><br>Red: forward stall site |

| | | |
|---|---|---|
| *Template2*<br><br>(*Templates3—6* are derived from *Template2* as described in the main text) | gtggtgaaaccgcatcttttgacatcgaagctaacggtaaagtgtacgtagaaaaaggccgcc<br>gtatcactgcgcgccacattcgccagctggaaaaagacgacgtcaaactgatcgaagtcccgg<br>ttgagtacatcgcaggtaaagtggttgctaaagactatattgatgagtctaccggcgagctgatct<br>gcgcagcgaacatggagctgagcctggatctgctggctaagctgagccagtctggtcacaag<br>cgtatcgaaacgctgttcaccaacgatctggatcacggcccatatatctctgaaaccttacgtgtc<br>gacccaactaacgaccgtctgagcgcactggtagaaatctaccgcatgatgcgccctggcga<br>gccgccgactcgtgaagcagctgaaagcctgttcgagaacctgttcttctccgaagaccgttat<br>gacttgtctgcggttggtcgtatgaagttcaaccgttctctgctgcgcgaagaaatcgaaggttcc<br>ggtatcctgagcaaagacgacatcattgatgttatgaaaaagctcatcgatatccgtaacggtaa<br>aggcgaagtcgatgatatcgaccacctcggcaaccgtcgtatccgttccgttggcgaaatggc<br>ggaaaaccagttccgcgttggcctggtacgtgtagagcgtgcggtgaaagagcgtctgtctctg<br>ggcgatctggataccctgatgccacaggatatgatcaacgccaagccgatttccgcagcagtg<br>aaagagttcttcggttccagccagctgtctcagtttatggaccagaacaacccgctgtctgagatt<br>acgcacaaacgtcgtatctccgcactcggcccaggcggtctgacccgtgaacgtgcaggcttc<br>gaagttcgagacgtacacccgactcactacggtcgcgtatgtccaatcgaaacccctgaaggt<br>ccgaacatcggtctgatcaactctctgtccgtgtacgcacagactaacgaatacggcttccttga<br>gactccgtatcgtaaagtgaccgacggtgttgtaactgacgaaattcactacctgtctgctatcga<br>agaaggcaactacgttatcgcccaggcgaactccaacttggatgaagaaggccacttcgtaga<br>agacctggtaacttgccgtagcaaaggcgaatccagcttgttcagccgcgaccaggttgactac<br>atggacgtatccacccagcaggtggtatccgtcggtgcgtccctgatcccgttcctggaacacg<br>atgacgccaaccgtgcattgatgggtgcgaacatgcaacgtcaggccgttccgactctgcgcg<br>ctgataagccgctggttggtactggtatggaacgtgctgttgccgttgactccggtgtaactgcg<br>gtagctaaacgtggtggtgtcgttcagtacgtggatgcttcccgtatcgttatcaaagttaacgaa<br>gacgagatgtatccgggtgaagcaggtatcgacatctacaacctgaccaaatacacccgttcta<br>accagaacacctgtatcaaccagatgccgtgtgtgtctctgggtgaaccggttgaacgtggcga<br>cgtgctggcagacggtccgtccaccgacctcggtgaactggcgcttggtcagaacatcgatga<br>ttatggcgaaactgaagtcagcgaaagggaagaaatttctctttggtttgttggcggttttcattatt<br>gcggcgtcggttgtgactcgcgcgaccatcggcggcgttatagaacagtacaatattccgctgt<br>ctgagtggacgacatcaatgtatgtgattcagtcatcgatgatttttgtttatagcctggtctttactg<br>tgttgctggcaatcccgttgggaatttatttccttggcggcgaagagcagtaa**GTAAAAA**<br>**ATAGGCCCGATAACTCGGGCCTTGTCAGTTATTGAAG**<br>**AGTCG**ttaatcgtcttcttcgtcatccagttcaacgggtgtctgatactggtcaggtttaatga<br>ccagcaggtcgcagcgaagatgatcaatcacctgttccgccgtgttgccgaggaatgctgctg<br>aaataccggtgcgtcctaccgtgcccagaaccacaatccccgcctgtaagtgctccgccaaat<br>caggaatcacctcttctggcagaccttttttctacgtgcgtcatgttttcattaatgccgaatttctgcc<br>gcagggctttcattgccagcaaatgttgcccacgaatggcatcgttataaacgctcgggtcaaat<br>tccggcagttcaatcgcgatattaattggcgttaccggataagcgccaaccagatgaacttcggt<br>atggttgacttgttctgccagttcgatcgtctctttgaccagtttttcattgagcgcattatgatacgg<br>ctcttcactggcgagattcaccgccaccagcgccttgcctccttccggccacggctggtctttca<br>ccatccacaccgggcttgggcatttgcgtaacagatgccagtccgttggcgtaaaaatcaccgc<br>ttccagacggtcatgttggtgcgccattttttagcaccaaatcgtgtccgccgctgatcacttcctg<br>aatgatggcttcgaaaggacggttatgccagaccactttaatttcaatgggaacgccagcattga<br>gataatattttgcctgctcgtgaatccaggctgtacgctggctgatgacgccctgacgcatagcg<br>gtacgttcgtccggggagagcagggtggtcatttcgtatgagaagtcatagatcggcaaaaag<br>gctttaattttgccaccaatccgttgatgtaaataaacagctcgccgcaatgctggttggtcgtcct<br>ggttaggatcgataacaacgagcatgttctgatacatagccatcagagtttagcgaatttttcgag<br>ggtgcgaataagctgtgtgacgaagccatattcgttatcgtaccaggcgaccgttttcaccagtt<br>gtaaatcgcccacggcggtaatttccgtttgcgtggcatcaaacaccgaaccgaaatggctgcc<br>aatgatatcggaagagactatttcttcatcggtataaccaaatgactcgttattggtggttgcttgttt<br>aagtgcgttattcacctcttcggcagtcactttttttccgagaatcgataccagttcagtgaccgaac<br>ctgttttcaccggcacgcgttgcgcatgacctttcagtttgccgctcagttccgggatcaccagac | Blue: forward repeat cassette<br><br>Gray: forward transcribed region<br><br>**Brown**: *ynaJ-uspE* bidirectional terminator<br><br>Black: reverse transcribed region<br><br>Magenta: reverse repeat cassette<br><br>Red: reverse stall site<br><br>Underline: reverse T7A1 promoter |

| | |
|---|---|
| | caatggcttttgccgcccccgtagtgtggggaatgatattttctgccgctgcgcgtgaagcacgt<br>aaatctttaccacgcgggccatccaccagtgactgggtgccagtataggcatgaatggtcgtca<br>tcgtgccgacttctatcccgaaactgtcatgcaaggctttggccatcggcgcaagacagttagtg<br>gtgcatgacgccacggaaacaatggtgtcgttgccatccagagtgtcgtcattgacgttataaac<br>gatagttttcatttcaccggcaggggcggaaatcaacaccttcttcgcaccagcatcaagatgcg<br>cctgcgatttctcggcggaggtataaaagccagtacattcgacaatgatttctgcacctttcgcttt<br>ccacggaatatttttagcctctttttcggcgtaaaccgcgatacttttcccatcaacgataagtgaat<br>cttccgtaaaatcaacgctccaggggaatggtccgtagtttgaatcatgtttcagcaggtaggcg<br>agaatttatggggaagtgagatcattaatagcgacaacgtctatgttgcttttgacttcaagtaatc<br>gacccaacaccagtcgaccgatacgaccaaaaccgttaataccaactttactcatggttttctcct<br>gtcaggaacgttcggatgagtttacggcgacggtcgatacgtacgaacaggttgcatgcgttaa<br>gataagggatagggagttaagataagggatagggagttaagataagggatagggagttaagat<br>aagggatagggagttaagataagggatagggagttaagataagggatagggagttaagataa<br>gggatagggGTGGATCCTTCATTCTCTCCATTCCACTTCAACT<br>CTCATATGGCTGTAAGTATCCTATAGGTTAGACTTTAAGT<br>CAATACTCTTTTTGATAAATTTTAAATTAATCTT |
